# A combinatorial mutational map of active non-native protein kinases by deep learning guided sequence design

**DOI:** 10.1101/2025.08.03.668353

**Authors:** Kosuke Seki, Amy B. Guo, Deniz Akpinaroglu, Tanja Kortemme

## Abstract

Mapping protein sequence-function landscapes has either been limited to small steps (only few mutations) or to sequences similar to those already explored by evolution to maintain activity. Here, we overcome both limitations by applying deep-learning guided redesign to a natural protein tyrosine kinase to generate novel, functional sequences with highly combinatorial mutations. Using cell-free assays, we measure the activities and concentrations of 537 redesigned sequences, which differ from the wild-type by an average of 37 mutations while retaining activity in 85% of variants. These sequences sample 436 unique mutations at 76 different positions throughout the kinase domain. A simple regression model identifies key sequence determinants of function and predicts the function of unseen sequences. Our approach demonstrates how integrating deep-learning guided redesign, functional measurement at scale, and interpretable computational modelling enables functional exploration of highly combinatorial and sparse sequence-function landscapes at mutational scales not possible before.

## Introduction

The ability to predict how amino acid sequences encode protein function - and how functions change with sequence variation - is important for both fundamental questions of protein evolution and for applied problems of optimizing proteins for desired properties. In principle, methods that efficiently sample large numbers of diverse yet functional protein sequences to generate high-quality training data could be used to build predictive models of protein function. Yet, such datasets and models that accurately predict function from sequence remain elusive. The effects of single mutations can be readily measured at scale^1,2^ and can often be predicted with useful accuracy.^3–5^ In contrast, exploring highly combinatorial sets of mutations is challenging because of scale and sparsity, since functional protein sequences are rare and most mutations are destabilizing.^6,7^

Due to these challenges, datasets that contain many quantitative activity measurements on combinatorially mutated yet functional sequences are rare. One way to overcome the sparsity of functional sequences is to measure the activities of sequence homologues of natural proteins, as they are constrained by evolution to be functional. Prior works have measured the activities of both natural enzyme sequences and designed sequences that recapitulate natural sequence covariation using a variety of methods, such as high-throughput *in vitro* assays or cellular selection assays.^8–10^ Other studies have traversed mutational pathways between protein sequences to explore their evolutionary pathways at scale.^11–14^ Yet, these approaches are inherently limited to the small fraction of sequence space that evolution has explored. Deep mutational scanning and directed evolution approaches can in principle explore novel sequences but in practice take relatively short mutational walks.^15,16^ *In vivo* hypermutation and continuous evolution approaches can bypass this constraint and identify functional sequences that highly diverge from their parent sequence, but only in the presence of an appropriate cellular selection assay.^17,18^ A generalizable method to efficiently explore the functional sequence space of any protein has thus been lacking, but would significantly improve the ability to map sequence-function relationships at scale.

Recent advances in deep learning models could now enable us to traverse rugged and massive sequence spaces more effectively. Structure-conditioned sequence design models, such as ProteinMPNN (pMPNN) and Frame2seq (F2s), among others, create sequences that fold into user-defined structures with high success rates.^19–21^ As sequences that fold into the correct structure are more likely to be functional, these sequence design models could be used to sample functional regions of highly combinatorial sequence spaces.^22^ Additionally, these models are capable of designing sequences distant from any natural counterparts. For example, F2s was used to design a protein with 0% identity to its native starting point, and the design was soluble, monomeric, and correctly folded into the target structure.^20^ Structure-conditioned sequence design models therefore could enable us to design sequences that are both functional and novel.

In this work, we apply structure-conditioned sequence design models to design thousands of diverse and new-to-nature protein tyrosine kinase domains (**Fig 1a**). We generate a dataset of 537 redesigned kinase sequences, each differing by an average of 37 mutations from the wild-type and by 30 mutations from each other. Using highly parallel, quantitative cell-free assays, we measure activity and expression levels of these variants, showing that 85% of them retain activity. We then use this dataset to train a ridge regression model that can accurately predict the specific activity of kinase variants unseen in training and can be interpreted to identify sequence determinants of function. Finally, we show that structure-conditioned sequence design models create sequences that deviate from nature. Our approach efficiently explores highly combinatorial sequence-function relationships by integrating deep learning guided sequence design, high-throughput quantitative measurement of functions, and simple interpretable modeling, and it lays the groundwork for exploring these relationships in other proteins at mutational scales not possible before.

**Figure 1:**
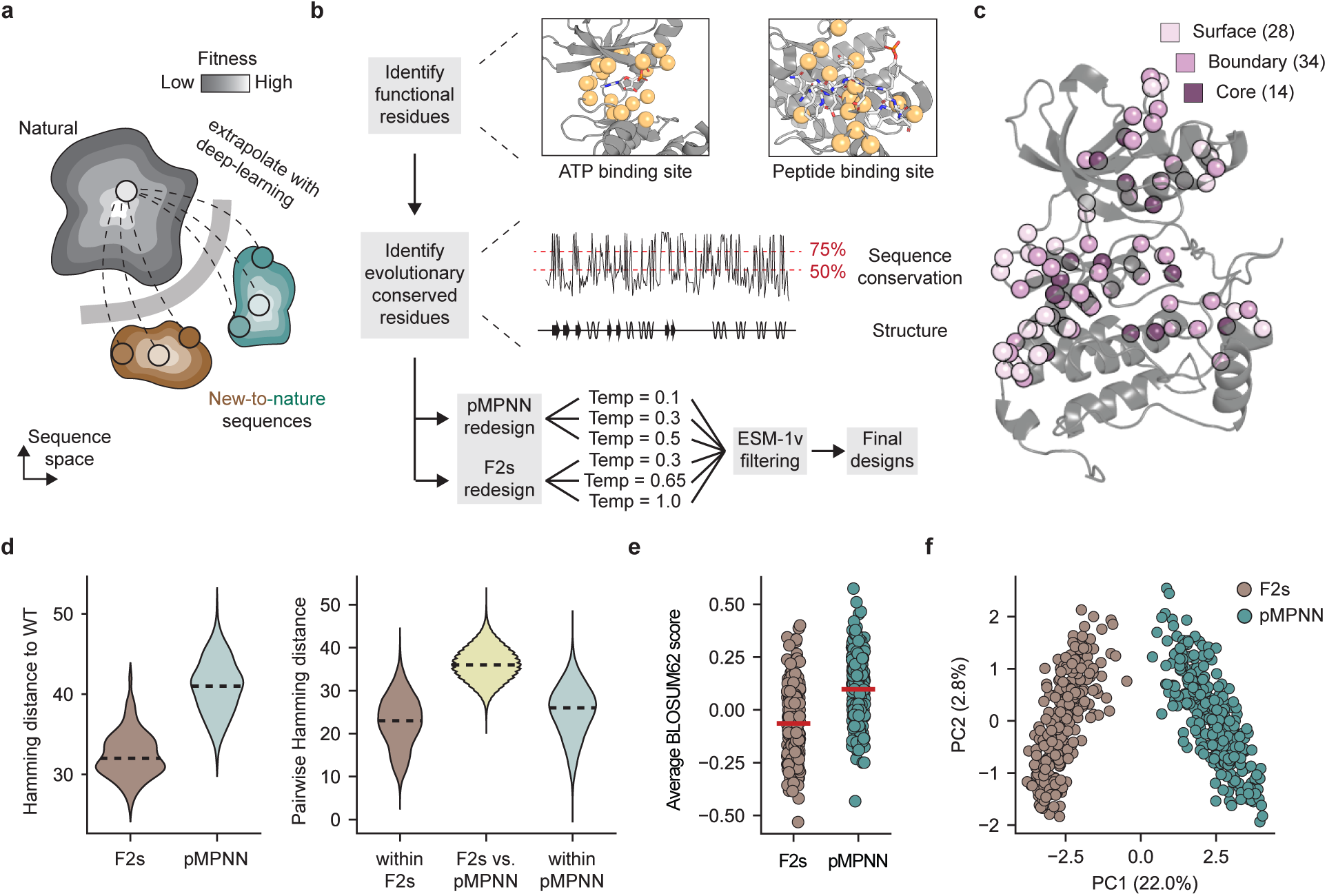
Deep learning models to guide efficient exploration of functional sequence space for EphB1. (**a**) Deep learning guided redesign of protein sequences to explore new-to-nature protein sequence spaces. (**b**) Workflow showing redesign strategy for EphB1. Functional and highly conserved residues are identified, sequences are redesigned using pMPNN and F2s with a range of sampling temperatures, and sequences are filtered using ESM-1v to enrich for functional sequences. (**c**) Redesigned positions plotted onto the structure of EphB1. Residues in the surface, boundary, and core regions are colored in shades of purple with numbers indicated in parentheses. (**d**) Distributions of Hamming distances between redesigned sequences and the reference (wild-type EphB1), as well as between and within sequences designed by the different deep learning methods, F2s and pMPNN. All redesigned sequences show large distance from the wild-type, and F2s sequences differ greatly from pMPNN sequences. Dotted lines show the mean Hamming distances. (**e**) Distributions of the average BLOSUM62 score of all mutations in each redesigned sequence. Red line indicates the mean BLOSUM62 score of all sequences within a category. (**f**) Principal component analysis of redesigned sequences shows that F2s and pMPNN sample distinct sequence spaces. PC1 discriminates between the two models.

## Results

### Pilot workflow and assay development

We first set out to develop an *Escherichia coli*-based cell-free expression (CFE) method to express and characterize receptor tyrosine kinase domains. CFE methods are compatible with high-throughput workflows,^8,23–26^ and can express toxic proteins such as receptor tyrosine kinases that would be challenging to heterologously express.^23,27^ To establish our experimental workflow, we selected the human EphB1 receptor tyrosine kinase domain (UniProt P54762, residues 602-896), which we refer to as EphB1, as a model system.^28^ The Eph receptors constitute the largest subfamily of receptor tyrosine kinases in humans and play key roles in intercellular signaling pathways.^29^ In the presence of protein folding chaperones, EphB1 was robustly expressed in the soluble fraction of CFE reactions (**Fig. S1a**). In addition, EphB1 activity was readily quantifiable in cell extracts without purification using PhosphoSens assays, which couple phosphorylation of a peptide substrate to fluorescence (**Fig. S1b**).^30^ We observed no background activity on the peptide substrate in the absence of the kinase (**Fig. S1b**). In contrast to CFE-based assays, we found that EphB1 is toxic to *E. coli* growth in the absence of a cognate phosphatase (**Fig. S2**). The toxicity and phosphatase activity would bias cell-based workflows, highlighting the advantage of our CFE-based method for EphB1 expression and testing.

We first designed a pilot library of EphB1 variants using the structure-conditioned sequence design model pMPNN (**Fig. 1b**).^19,31^ We fixed EphB1 residues containing a Cβ within 7 Å of ATP in a representative crystal structure (PDB ID: 7kpm) and used a structural alignment to the EphB1 homolog EphA3 (PDB ID: 3fxx) to identify and fix peptide binding residues. To identify evolutionarily conserved residues, we built multiple sequence alignments (MSAs) of natural tyrosine kinase domains from InterPro (ID: IPR020635) and curated three sets of evolutionarily conserved residues using conservation thresholds of 50, 75, and 95%. All residues under these thresholds were allowed to be mutated (“redesignable”) by pMPNN (**Methods**). In parallel, we created a fourth residue set where all core and boundary residues were fixed to assess the effect of only redesigning the EphB1 surface. 300 sequences for each of the 4 sets of fixed residues were designed using both vanilla pMPNN and soluble pMPNN,^32^ and we selected three sequences within each set corresponding to sequences with the maximum ESM-1v score, the minimum ESM-1v score, and the minimum F2s score for experimental testing (**Table S1**). These scores have been shown to be useful filtering metrics to capture sequence and structure information (**Fig. S3**).^33,34^ We also predicted the structure for every sequence using AlphaFold2 in single-sequence mode, and selected one additional sequence with the highest AF2 confidence metric pLDDT of 79.6 (**Methods**). In total, the selected 25 variants contain 68 – 114 mutations from the wild-type EphB1 kinase domain.

We used CFE-based assays to rapidly characterize these variants. Compared to the wild-type, almost all redesigned EphB1 variants showed significantly improved expression and solubility, particularly in the absence of chaperones (**Fig. S4**). In PhosphoSens assays, five redesigned EphB1 variants show considerable activity (> 1.5x) compared to the no enzyme control while the remaining twenty were nonfunctional (**Fig. S5**, **Table S1**). We define activity in PhosphoSens assays here as the phosphorylation rate measured by the increase in fluorescence over time from dilute CFE reactions, which reflects both enzyme expression and catalysis. Of these five functional variants, three were designed with a conservation threshold of 50% and two with a threshold of 75% (**Fig. S5**). Three of the active sequences were selected based on having a maximum ESM-1v score, and the remaining two were selected based on the F2s score (**Table S1**). The best redesigned variant, Sol50-0.3-89, had 87 mutations and activity comparable with the wild-type enzyme. These results show that structure-conditioned sequence design models can make large jumps in sequence space with notable success rates even when only testing a few variants.

### Second generation EphB1 library and sequence diversity

We next designed a second-generation library of EphB1 variants to explore sequence function relationships at a larger scale. Because of size limitations in pooled DNA synthesis, we chose to redesign a 150-residue long region of EphB1 (residues 661-811) which encompasses important functional regions such as the αC helix, ATP binding site, the catalytic loop, and the activation loop (**Fig. 1c**). We also explored slight modifications to the redesign strategy used for our pilot library. Residues under the 50% conservation threshold were always redesignable, but we also randomly selected a subset of residues between 50 - 75% conservation to be redesigned in each round of sequence design. Each round of redesign in the second-generation library therefore uses different sets of fixed residues. We expected that this strategy would increase sequence diversity without compromising function. Second, we used F2s to design a second set of sequences to circumvent a library bias that could result from using only a single sequence design model.^20^ In total, we designed 60,000 sequences in 100 rounds of sequence design, scored the sequences using ESM-1v, and included the top 1,000 sequences from each pMPNN and F2s for experimental testing within a pooled DNA library of 2,000 sequences (**Methods**, **Fig. S6**).

The redesigned EphB1 library encompasses a broad subset of sequence space. In 537 experimentally tested sequences of this library (discussed in the next section), the designs contain an average of 37 mutations relative to the wild-type at 76 different positions (**Fig. 1c,d, Fig. S7**). This corresponds to a theoretical complete combinatorial space of 5.4 x 10^46^ sequences (**Methods**, **Table S2, Fig. S7**). The tested subset of sequences samples 436 of 570 unique mutations in the total library, and these mutations are structurally distributed throughout the core, boundary, and surface of EphB1 (**Fig. 1c**). We find that using both pMPNN and F2s adds to the sequence diversity of the library. pMPNN and F2s made an average of 40 and 32 mutations to the wild-type, respectively (**Fig. 1d**). The average pairwise Hamming distance within tested pMPNN and F2s sequences is 25 ± 6.3 and 23 ± 6.2, respectively, and increases to 36 ± 4.0 between pMPNN and F2s sequences (**Fig. 1d**). Substitution matrices of pMPNN and F2s sequences show the different mutational preferences of each model (**Fig. S8**), and average BLOSUM62 scores of each sequence show that F2s incorporates slightly less conservative mutations (p = 1.26 e-27 by Mann-Whitney U test, **Fig. 1e**). To identify the mutational signatures of each model, we conducted principal component analysis (PCA) and found that PC1 discriminates between sequences designed by each model (**Fig. 1f**). Loadings onto PC1 reveal mutational signatures of F2s and pMPNN, which typically redesign similar positions but incorporate different amino acids (**Fig. S9**). The complementary use of both F2s and pMPNN therefore enables broader exploration of sequence space compared to using only a single model.

### Experimental evaluation of redesigned EphB1 sequences at scale

We developed a semi-automated, CFE-based workflow to biochemically characterize redesigned sequences at scale using robotic colony pickers and liquid handlers (**Fig. 2a**).^35^ We cloned a pooled plasmid library of all 2,000 redesigned EphB1 variants and confirmed coverage of each library member using Next Generation Sequencing (NGS). We prepared arrayed linear DNA templates compatible for CFE by transforming the library into *E. coli* at > 10x coverage, picking individual colonies, growing cell cultures, and amplifying DNA templates directly from *E. coli* cultures using PCR. The redesigned region within each DNA template was barcoded and sequenced using NGS to identify each library member. Finally, we expressed and measured the activity of redesigned EphB1 variants using CFE. To quantify specific activity levels, we normalize phosphorylation activity measured in PhophoSens assays using expression levels measured by complementation of split luciferase with an N-terminal HiBiT tag in each kinase variant.^36,37^

**Figure 2:**
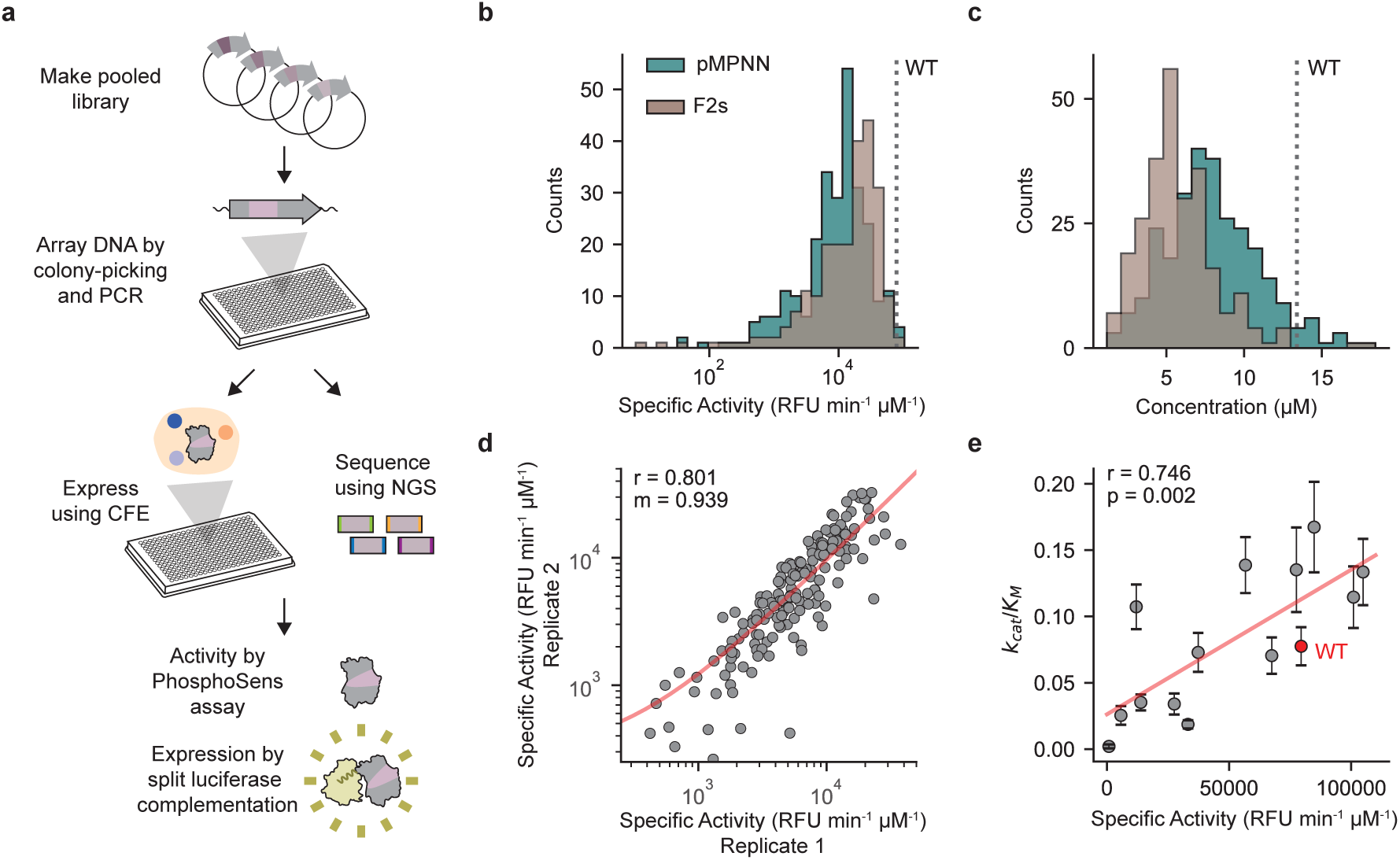
Quantitative activity measurements of hundreds of redesigned EphB1 variants. (**a**) A CFE-based workflow enables performing thousands of reactions in parallel. Redesigned kinase variants are characterized by PhosphoSens assays to measure activity and split luciferase complementation to measure expression. (**b**) Histograms of specific activities of all kinase variants. F2s sequences are generally more active than pMPNN sequences. (**c**) Histograms of expression levels of all kinase variants. pMPNN sequences are generally more highly expressed. Wildtype levels in (b) and (c) are marked with a dotted line. (**d**) Scatter plots of replicate measurements between different days show strong reproducibility (Pearson r: 0.801, m: 0.939). Points represent measurements of a single sequence between different days, and the fit to replicate measurements is shown in red. (**e**) Scatter plot between plate-based measurements of specific activity in cell extracts and Michaelis-Menten parameters with individually purified proteins, showing a strong correlation between the two measurements (Pearson r = 0.746, p = 0.002). Points represent the average from n = 3 replicates and error bars show one standard deviation of the fitted parameters.

We measured the activities of hundreds of redesigned variants using this workflow. Within a single production run, we picked 1,536 colonies, assigned single sequences to 955 wells, measured activity and expression for 715 unique sequences, and filtered for 537 sequence-function measurements that pass quality control metrics (**Fig. S10**). The sequence space we sample is highly enriched for function; out of 715 unique sequences, 605 (85%, **Fig. S10**) have measurable activity (> 1.5x the negative control) despite large divergence from the wild-type sequence (**Fig. 1d**). Specific activities range from 0 – 1.3x wild-type activity levels (**Fig. 2b**) and expression levels range from 0 – 1.4x wild-type expression levels (**Fig. 2c**). Finally, the workflow is robust; day-to-day replicate measurements of a 384-well plate are highly correlated and show little variation (Pearson r = 0.80, m = 0.939, **Fig. 2d**), and internal replicates of sequences (n = 92 duplicates, n = 13 triplicates) that occur due to random sampling of colonies are highly correlated (Pearson r = 0.786, **Fig. S11**).

Sequences designed by F2s and pMPNN have distinct functional properties. F2s sequences are significantly more active compared to those designed by pMPNN (p = 2.83 e-3 by Mann-Whitney U test) but show worse expression (p = 3.03 e-19 by Mann-Whitney U test) (**Fig. 2b,c**). The improved specific activity of F2s sequences might result from a shorter Hamming distance to the wild-type, as specific activity decreases with added mutations for both models, despite the less conservative mutations that are generally sampled by F2s (**Fig. S12, 1d,e**). We observe a slight tradeoff between specific activities and expression levels in both F2s and pMPNN sequences, with variants with high specific activity displaying overall lower expression levels and variants with high expression levels having moderate specific activities (**Fig. S13**).

Specific activity measurements from CFE assays were confirmed to reflect catalytic parameters by conducting Michaelis-Menten analysis for 13 individually purified redesigned EphB1 variants (**Table S3**, **Fig. 2e**). All redesigned variants are soluble and monomeric as assessed by size exclusion chromatography (**Fig. S14**). Michaelis-Menten curves are well fit to the data and k_cat_/K_M_ values are well correlated with measurements from plate-based assays (Pearson r = 0.746, **Fig. S15, 2e**). Specific activity measurements are correlated to k_cat_ rather than K_M_, which is expected as putative peptide binding residues were fixed (**Fig. S16**). The best pMPNN and F2s variants from this analysis have k_cat_/K_M_ values that are 2.14x and 1.72x greater than that of the wild-type, respectively. Our redesigned variants thus sample functional sequences that can exceed the activity of the reference natural kinase domain as well as those that span a range of attenuated activity levels. In total, these results show how our assays can capture biochemical parameters at high throughput to explore functional sequences at large scales.

### Building a predictive model

Using this high-quality dataset of labelled sequences, we next aimed to train models that accurately predict the function of unseen sequences. As an initial test, we assessed if principal component analysis could uncover any patterns of activity within our redesigned sequences. We found that principal components of neither one-hot encoded (OHE) sequences nor ESM-1v embeddings capture activity (**Fig. S17**). In both cases, the largest (but weak) correlation is from the first principal component (ESM Pearson r = 0.252, OHE Pearson r = -0.184), which likely only captures differences between F2s and pMPNN sequences (**Fig. 1f, S17**).

Previous studies have shown that simple supervised machine-learning models can be effective predictors for activity, but these models have typically only been evaluated on sequences containing 1 - 4 mutations or mutations at up to 10 positions.^26,38,39^ We trained simple models (ridge regression, lasso regression, and random forest) on labelled sequence features (OHE, ESM-1v, Georgiev)^34,40^, combined with either evolutionary information (EVcouplings)^41^ or predicted changes in protein stability (ΔΔG) compared to wild-type (ThermoMPNN).^3^ As our measured specific activities were right-skewed, we transformed our data towards normality by applying a Yeo-Johnson transformation to train our models (**Fig. S18**). We report the average Mean Squared Error (MSE) and Spearman correlation coefficient (r) across the test folds in 5-fold cross validation with randomly selected training and test splits (**Fig. 3a**). Even with diverse sequences, we find that simple models have strong performance on our dataset. The best performing model is ridge regression with ESM-1v embeddings combined with EVcouplings information (Spearman r = 0.756, MSE = 420 arbitrary units, a.u.). Including zero-shot predictions of mutation effects from EVcouplings or ThermoMPNN as additional features does not substantially improve performance compared to only using ESM-1v embeddings (Spearman r = 0.755, MSE = 422 a.u.). This result is not surprising, as ESM-1v embeddings likely capture evolutionary and structural information.^42^ Consistent with this reasoning, we find that ESM-1v embeddings typically perform slightly better than residue level features (OHE, Georgiev). In total, our data show how simple machine learning models can perform well on sequences with highly combinatorial mutations.

**Figure 3:**
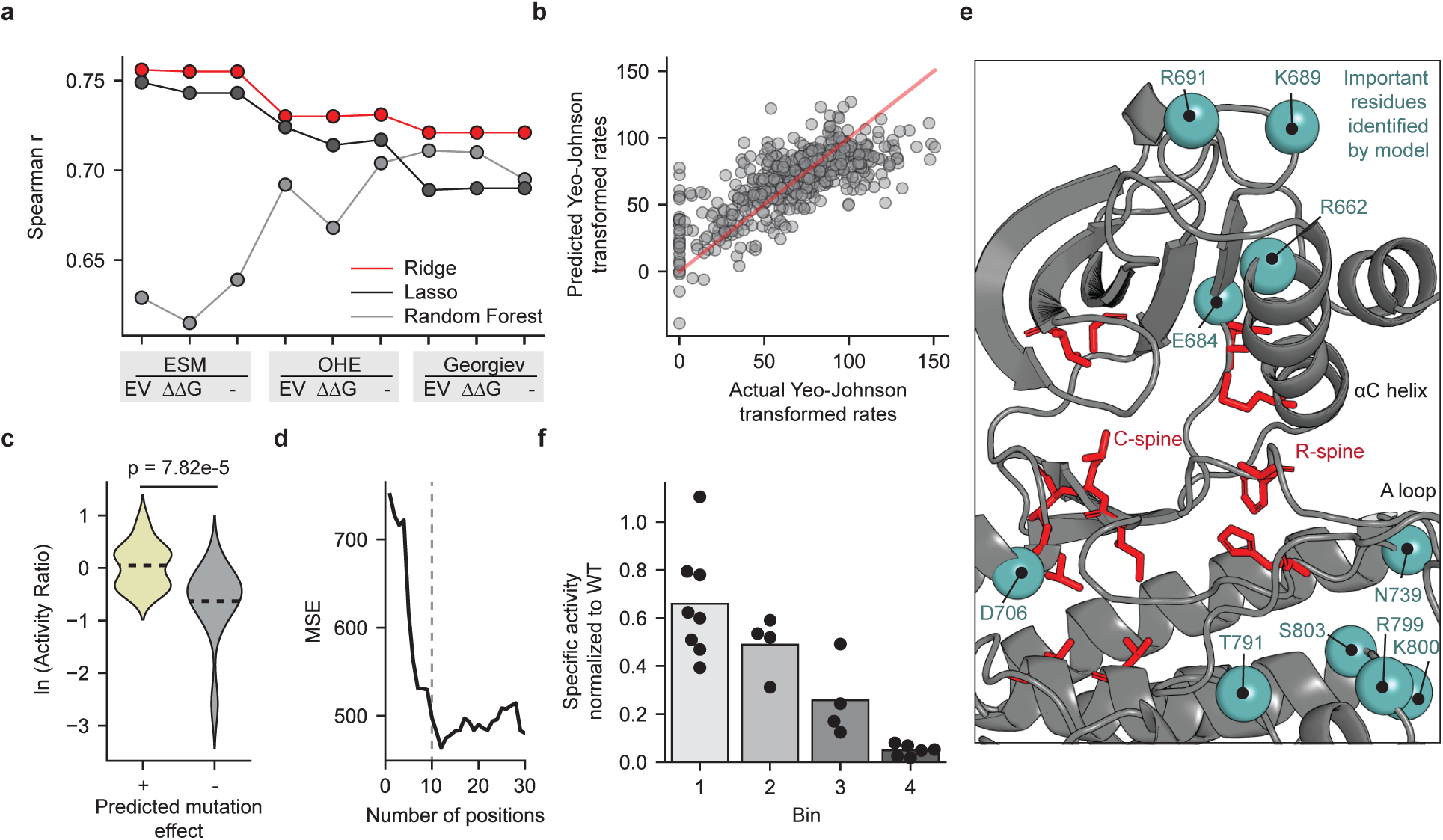
Simple machine learning models accurately learn sequence-function relationships. (**a**) Spearman correlation coefficient of predicted to measured specific activities for different simple machine-learning models using a combination of sequence features (ESM-1v embeddings, OHE, or Georgiev physiochemical encodings) augmented with zero-shot predictors (EV: EVCouplings, ΔΔG: ThermoMPNN predictions, -: no augmentation). Ridge regression models perform best. (**b**) Scatter plot of experimentally determined rates (transformed by Yeo-Johnson) for all kinase variants versus rates predicted by ridge regression models trained on one hot encoded sequences. Points show the predicted activities from sequences in the test folds during 5-fold cross validation with random splits. Linear best fit is shown as red line (Spearman r = 0.731). (**c**) Distributions of the fold-change between the average specific activities of sequences with and without a given mutation of interest. Only mutations where the distributions of sequences with and without the mutation of interest were statistically different (n = 49, adjusted p < 0.05 by Mann-Whitney U test after Benjamini/Hochberg correction) are shown. The predicted effects of mutations (given by the sign of the regression coefficient) generally agrees with experimental observations. Dotted line shows the average fold-change of mutations with positive or negative regression coefficients. (**d**) Dependence of MSE on number of positions included in model training. The model depends only on ∼10 positions to accurately predict activity. Each position is rank ordered based on the sum of the absolute values of regression coefficients at that position. Dotted line is x = 10 positions. (**e**) Important positions identified by the model plotted on EphB1 structure, labelled as teal spheres, are near known regulatory elements of protein tyrosine kinases. Residues in the spines are labelled in red. (**f**) Bar plot of experimental measurements of specific activity of sequences unseen by the ridge regression model trained on one-hot encoded sequences. Specific activities were normalized to the specific activity of the wild-type measured in the same experiment. The magnitudes of specific activities are qualitatively consistent with model predictions. Each point reflects the average specific activity of n = 3 replicates on a single variant, the bars represent the average specific activity of variants within each bin, and the bins are rank ordered by predicted activity (1: top 8, 2: high, 3: medium, 4: low/inactive).

The simple architecture of these models enables us to identify sequence determinants of kinase function. To facilitate biophysical interpretation, we trained a ridge regression model only on OHE sequences as input features (Spearman r = 0.731, MSE = 468, **Fig. 3a,b**). First, we analyzed the regression coefficients to learn the general impact of each residue on function (**Fig. S19**). The model identifies biologically important functional residues. For example, regression coefficients for mutations to positions T791 and R799, which are conserved within functional members of the human Eph receptor tyrosine kinase subfamily but not within non-functional pseudokinases EphA10 and EphB6, are strongly negative.^43^ The regression coefficients also broadly reflect the effects of individual mutations in different sequence contexts. The average activity of sequences containing R662E, a mutation with a strong positive coefficient, is 2.1x greater than those without R662E (p = 1.57 e-8 by Mann-Whitney U test, **Fig. S20a**). Similarly, sequences with K800D, a mutation with a strong negative coefficient, are 0.51x as active as those without it (p = 1.93 e-8 by Mann-Whitney U test, **Fig. S20b**). In general, sequences containing predicted negative mutations are, on average, 0.531x as active as sequences without those mutations, while sequences with positive mutations display a modest 1.05x increase in activity (n = 49 statistically significant differences, **Fig. 3c**). The broad distribution of these functional effects is unsurprising, as the consequences of these mutations could change depending on their specific sequence contexts (**Fig. 3c**). This may reflect effects from epistasis or from stronger functional effects from other mutations.

To identify the importance of each position to model performance, we next trained models that use limited subsets of residues to predict activity (**Fig. 3d**). Strikingly, model performance when using sequence information at ∼10 positions is comparable to that when using all available sequence information (T791, D706, R691, S803, R799, R662, N739, E684, K800, K689, in order of most importance, i.e. the sum of the magnitude of all ridge regression coefficients at a given position). The Shannon entropies of mutation distributions at these positions vary between 0.521 and 2.27 bits, suggesting that the model is not focusing only on positions with high variance (**Fig. S21**). Consistent with the interpretation that the model is identifying the relative importance of each residue and mutation, many mutations at these positions appear to converge in sequences with specific activity > 15,000 RFU min^-1^ µM^-1^ (**Fig. S22a**). Nearly all active sequences converge on T791, the highest ranked position. We also observe strong enrichment (defined as the difference in mutation frequences between active and all sequence variants) of R662E and R691K in addition to strong depletion of K800D and the wild-type D706 relative to their starting frequencies (**Fig. S22a**). Interestingly, some of these mutations are rare in nature (**Fig. S22b**); E is found at position 662 in only 3.52% of EphB1 homologues and 1.91% of representative natural protein tyrosine kinases. This suggests that the model can identify beneficial mutations even when they are rare in nature. Structurally, these 10 positions are clustered around known regulatory regions of the EphB1 kinase (**Fig. 3e**, **S23**). Seven of these positions are within 7 Å of functional regions, such as the R spine, C spine, ATP binding pocket, peptide binding pocket, αC helix, and activation loop (**Fig. S23**). These regions are known to regulate catalytic activity in all protein kinases; for example, formation of the R-spine stabilizes the catalytic conformation of protein kinases.^44,45^ Mutations near these regions may therefore play functional roles in catalysis. Taken together, these data show how simple models can learn features of complex sequence-function relationships.

We then asked if the model could accurately predict specific activities of variants outside of the 537 sequences in its training set. We predicted the activity of 1,228 unseen sequences within our 2,000-member library and selected 22 sequences across four bins of predicted activity for experimental validation (Bin 1: top 8, Bin 2: high, Bin 3: medium, Bin 4: low/inactive, **Table S4**). On average, these sequences are 13 mutations away from their closest relative in the training data and contain 6.8 mutations in the top 10 most important residues in the regression model (**Table S4**). In experimental measurements, we find that the average specific activity within each bin closely aligns with model predictions (**Fig. 3f**). The model therefore can qualitatively predict kinase activity from sequence alone and extrapolates to sequences outside of its training dataset.

### Sequence novelty

Lastly, we analyzed the divergence and novelty of redesigned EphB1 variants. One possibility is that the structure-conditioned sequence design models simply recapitulated natural sequence variation found within the Eph receptor tyrosine kinase family. To probe this question, we calculated the statistical energy of each redesigned sequence using a Potts model trained on EphB1 homologues.^46^ The trained Potts model serves as a suitable proxy for sequence variation in natural sequences, as it is trained on Eph receptor homologues and recapitulates observed structural contacts in EphB1 (**Fig. S24**). The Potts model should score sequences poorly if they have unnatural sequence patterns. Compared to natural Eph receptor tyrosine kinases, redesigned sequences score systematically worse (higher energy, p = 7.97 e-89 by Mann-Whitney U test, **Fig. 4a**). This finding suggests that structure-based sequence design methods are sampling mutations that deviate from natural variation in homologues. In addition, we find that the statistical energy only has a weak correlation with measured specific activities (Pearson r = - 0.285, **Fig. 4b**). Sequences that score poorly in Potts models can still retain high activity (**Fig. 4b**). As a simpler regression model can predict function well (**Fig. 3a,b**), this suggests that our redesigned sequences are likely out-of-distribution of the natural sequences used to train the Potts model.

**Figure 4:**
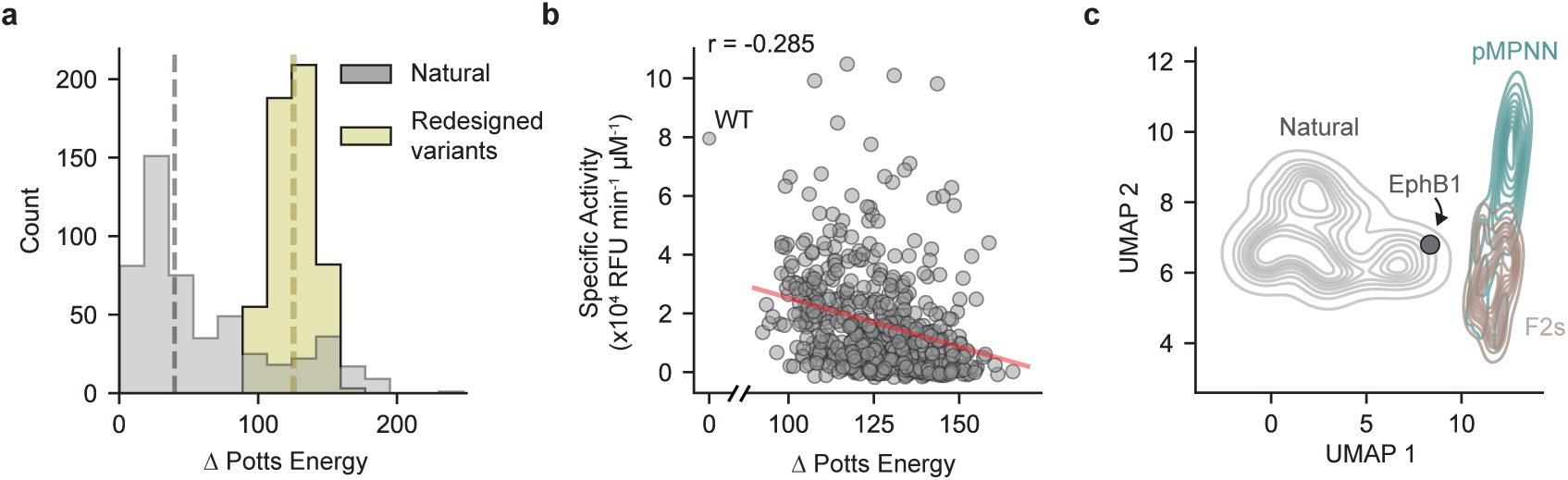
Redesigned EphB1 variants deviate from natural sequences. (**a**) Distributions of statistical energies of redesigned EphB1 variants and natural Eph receptor tyrosine kinase domains calculated by Potts models relative to EphB1. Redesigned EphB1 sequences systematically score higher (worse) than natural Eph receptor tyrosine kinase domains by a Potts model (p = 7.97 e-89 by Mann-Whitney U test). Dotted lines show the median Potts energies for natural and redesigned variants, respectively. (**b**) Scatter plot of statistical energies calculated by a Potts model for redesigned EphB1 variants versus specific activities, showing only a weak correlation (Pearson r: -0.285). Linear regression fit is shown as a red line. (**c**) UMAP visualization of ESM2 embeddings of active EphB1 variants, designed with F2s and pMPNN, and natural kinase sequences. Designed kinases cluster separately from representative natural tyrosine kinase domains and from the reference wild-type EphB1 sequence.

As a second proxy for divergence of redesigned variants from natural sequences, we visualized ESM2 embeddings, as embeddings from state-of-the-art protein language models capture evolutionary information from natural sequences. We retrieved representative sequences from all natural protein tyrosine kinase domains by clustering, generated ESM2 embeddings for each sequence, and visualized the embedding space using UMAPs (**Fig. 4c**). We observe that pMPNN and F2s designed EphB1 sequences that cluster somewhat separately from each other but entirely separate from natural protein tyrosine kinase domains. The wild-type sequence clusters with natural ones. This finding suggests that structure-based sequence design models are deviating not only from natural Eph receptor tyrosine kinases, but also from natural tyrosine kinase domains generally. Taken together, these data suggest that deep learning-based sequence design models can generate sequences distant from nature.

## Discussion and conclusion

Our work shows how functional protein sequence landscapes can be efficiently traversed and measured using a combination of deep learning-guided redesign and high-throughput quantitative assays. Deep learning-guided protein design methods enabled us to explore functional sequence spaces with high success rates.^20^ Highly parallel cell-free assays allowed us to quantitatively characterize the activities of 537 redesigned EphB1 variants and train simple supervised machine learning models that both reveal sequence-level determinants of function and accurately predict the activity of unseen variants. For example, our model identified T791 as a key position determining kinase activity, and a recent deep mutational scan of the Src kinase found that all mutations to the equivalent position, T432, were deleterious.^47^ More broadly, the dataset we generate explores a broader sequence-function landscape compared to other works,^48^ as our sequences contain an average of 37 mutations across 76 positions. We expect that this dataset will be valuable for evaluating and training protein function prediction models.

The theoretical sequence space (10^46^) for the mutations we sample is astronomically larger than the number of sequences we generated data for. While advances in deep learning models and scoring methods were critical in focusing our experimental work on sequences that were likely to be functional, our sampled subset will not represent all functional sequence space. Advances in structure-conditioned sequence design models that integrate ligand-level and atomic-level information might enable more diverse exploration of functional sequence space.^49,50^ In addition, while our experimental approach can identify key determinants of function (such as T791 in EphB1), identifying higher order interactions that are responsible for phenomena such as epistasis or allostery may require significantly more data.^13,51^ Methodological advances to measure protein function at large scales, such as microfluidics-assisted assays or *in vitro* display, will be needed for functions that require a large, quantitative dataset.^2,52^

In principle, our approach could create and mine large and diverse sequence-function datasets for protein functions that can be measured in high-throughput using cell-free platforms. As this platform can be readily adapted to many assays compatible with microtiter plates, it should be generalizable to many different types of functions. Extensions of this platform within protein tyrosine kinases could be used to either measure other desirable properties, such as substrate specificity, or engineer ones not found in natural tyrosine kinases. Further, as many protein functions have now become designable with deep-learning approaches, we expect that our workflow could be useful for dissecting sequence-function landscapes of other *de novo* designed enzymes.^53–55^ This approach should therefore have broad utility in mapping the sequence-function landscapes of proteins, identifying the key sequence determinants of function, and engineering new-to-nature enzymes with tailored properties.

## Methods

### Experimental Methods

#### Materials

All chemical reagents were purchased from Sigma-Aldrich unless otherwise noted. All primers were purchased from Integrated DNA Technologies. Standard antibiotic concentrations were 50 µg/mL for Kanamycin (Kan), 50 µg/mL for Streptomycin (Strep), and 34 µg/mL for Chloramphenicol (Cm). Minipreps were performed using Qiagen QIAprep Spin Miniprep Kits.

#### DNA amplification

PCR was performed using Q5 2x Master Mix (New England Biolabs, NEB). DNA templates for PCR were either 1 ng of gene fragments (Twist Biosciences), 1 ng of miniprepped plasmids (Twist Biosciences), or 10% v/v of saturated cell culture. Primer sequences for all amplifications are listed in supplementary files. Thermocycling parameters are listed below. T_m_s (step 3 in table below) were calculated for each primer set using the NEB T_M_ calculator (https://tmcalculator.neb.com/#!/main) and was optionally optimized with a temperature gradient from 72 – 55 °C. Extension times were calculated for each amplicon using the recommended 25 s / kb. PCR amplification was confirmed using agarose gel electrophoresis. 2 µL of PCR was mixed with 1 µL of 6X Loading dye (NEB) and 3 µL of water. Samples were loaded into a 1% w/v agarose gel made with SYBR Safe (ThermoFisher) and run at 120V for 30 – 45 min. Gels were imaged in a Bio-Rad Gel-Doc EZ imager.

| Step | Temperature (°C) | Time (s) |
| --- | --- | --- |
| 1 | 98 | 30 |
| 2 | 98 | 10 |
| 3 | Variable | 30 |
| 4 | 72 | Variable |
| Repeat 2-4 34X |  |  |
| 5 | 72 | 300 |
| 6 | 12 | Hold |

#### DNA cloning

General protocols are described with specific details below. All cloning was done using Gibson assembly with NEB HiFi DNA Assembly Master Mix or an in-house Gibson assembly mixture.^56^ Linear DNA appropriate for Gibson assembly was amplified using Q5 PCR. For PCRs with plasmid templates, PCRs were purified by PCR cleanup kits (Qiagen) in water and digested in 1X CutSmart Buffer with Dpn1 at 37 °C for at least 2 hours. Digested PCR products were used without purification for Gibson Assembly. In general, ∼ 10 ng of backbone was mixed with a 3X molar excess of insert, 5 µL of 2X HiFi Master Mix, and water to 10 µL according to manufacturer protocols. Reactions were incubated at 50 °C for 30 min – 1 hr. NEB 5α cells were transformed with 3 µL of Gibson assembly reaction according to manufacturer protocols. After recovery, the entire culture was spun down at 18 k x g for 30 seconds, resuspended in 100 µL of LB, and streaked onto LB-agar plates with appropriate antibiotics. Plates were incubated at 37 °C overnight. Five single colonies were inoculated into 5 mL of LB with appropriate antibiotics. Plasmid DNA was purified by miniprep and sequence-verified either by whole plasmid sequencing (Plasmidsaurus) or Sanger sequencing (Elim).

pCDF-PTP1b: A pCDF-PTP1b (Strep) plasmid was created by Gibson assembly, as described above, with a PTP1b gene fragment (Twist Biosciences) and a linearized pCDF backbone (Novagen). The pCDF backbone was linearized by PCR using primers pr3 and pr4. The pCDF backbone was gel purified due to formation of non-specific products using Zymo Gel DNA Recovery kits. Plasmids were assembled and isolated as described above.

pJL1-EphB1: All cloning steps with pJL1-EphB1 required co-transformation of pCDF-PTP1b to prevent mutations to EphB1. pJL1 backbone was prepared by PCR using pr77 and pr78 on the pJL1-sfGFP plasmid (https://www.addgene.org/102634/) and was assembled with an N-terminal His-tag-EphB1 gene fragment (Twist Biosciences) that was amplified by PCR using pr79 and pr80. Plasmids were assembled and isolated as described above. The N-terminal His-tag was exchanged for a HiBiT tag using an intramolecular Gibson assembly of the PCR product using pr105 and pr106 on the resulting plasmid. Plasmids were isolated as described above. Plasmids were sequence confirmed by lifting the open reading frame of pJL1 using PCR with pr85 and pr87 and Sanger sequencing using T7 forward and T7 reverse primers.

pET-EphB1 (and variants thereof): All cloning steps with pET-EphB1 required co-transformation of pCDF-PTP1b to prevent mutations. A pET-EphB1 (Kan) plasmid containing a TEV-cleavable N-terminal His tag was purchased from Twist Biosciences. Gibson assembly, as described previously, was used to clone all EphB1 variants into this backbone. To clone Sol50-0.3-89 into this pET backbone (**Table S1**), a linearized pET backbone was synthesized using PCR with primers pr81 and pr82 and assembled with a linearized Sol50-0.3-89 gene made by PCR using primers pr83 and pr84 from an eBlock (IDT). To clone variants with a redesigned fragment (**Table S2**), a pET backbone was linearized by PCR using primers pr171 and pr172 and combined with a linearized EphB1 variant fragment made by PCR using primers pr169 and pr170 from resuspended pellets of cell cultures. All plasmids were isolated as described above. Plasmids were sequence-verified by using pr79 and pr80 to amplify the open reading frame of the pET plasmid and Sanger sequencing using pr169 and pr170 to verify the redesigned fragment.

#### S30 E. coli lysate preparation

BL21 Star (DE3) (Invitrogen) was used as the chassis strain for all S30 lysates. To enrich S30 lysates with chaperones, BL21 Star (DE3) was transformed with pG-KJE8 (Takara Bio, Cm resistance). Cm was used in subsequent steps as needed. Cells were streaked onto LB-agar plates with appropriate antibiotics. A single colony was inoculated into 5 mL of LB media with appropriate antibiotics and grown at 37 °C overnight with 220 rpm shaking. Overnight cultures were inoculated into 1L of 2xYTPG (16 g/L tryptone, 10 g/L yeast extract, 5 g/L NaCl, 7 g/L potassium phosphate monobasic, 3 g/L potassium phosphate dibasic, and 18 g/L glucose) with appropriate antibiotics in 2.5 L TunAir flasks (IBI Scientific) at OD_600_ = 0.075. Cultures were grown at 37 °C with 220 rpm shaking. Cultures were induced with 1 mM IPTG at OD_600_ = 0.6 to express T7 RNA polymerase. To induce chaperone expression, 1 g of arabinose was added at the same time. At OD_600_ = 3.0, cells were pelleted by centrifugation at 5k x g for 15 min at 4 °C. Cells were washed three times by resuspending cells in 25 mL of S30 buffer (10 mM tris acetate pH 8.2, 14 mM magnesium acetate, and 60 mM potassium acetate) by vortexing, pelleting cells by centrifugation at 3.5 k x g for 5 min at 4 °C, and decanting supernatant. Tubes were thoroughly cleaned to remove residual S30 buffer. Cells were stored at -80 °C or processed immediately into lysate.

Cells were resuspended in 1 mL of S30 buffer per gram of wet cell mass by vortexing. 1 mL of cell suspension was aliquoted into 1.5 mL Eppendorf tubes. Cells were lysed by sonication for 45 s using 5s on / 5s off cycles at 15% amplitude using a Sonic Dismembrator Model 500 (Fisher). The sonication time was optimized for best yields in CFE reactions. Lysates were centrifuged at 12k x g for 10 min at 4 °C. The soluble fraction of the cell lysate was consolidated, split into 50 µL and 500 µL aliquots, and flash frozen as single use aliquots in a dry ice ethanol bath. S30 lysates were stored at -80 °C.

#### Cell-free expression

Cell-free expression reactions are based on the previously published PANOX-SP system.^57^ Reactions consisted of 8-12 mM magnesium glutamate, 10 mM ammonium acetate, 0.130 M potassium glutamate, 1.2 mM ATP, 0.85 mM GTP, 0.85 mM UTP, 0.85 mM CTP, 0.03 mg/mL folinic acid, 0.17 mg/mL tRNA, 0.4 mM NAD, 0.27 mM CoA, 4 mM oxalic acid, 1 mM putrescine, 1.5 mM spermidine, 57 mM HEPES pH 7.2, 2.5 mM of each amino acid, 33 mM PEP, 30% v/v S30 *E. coli* extract, and 13% v/v µL DNA. The optimum concentration of magnesium glutamate was tested for every batch of S30 *E. coli* cell extract and ranged between 8 -12 mM. Unpurified linear PCR products were used as DNA templates for all CFE reactions. Linear templates for CFE were prepared using the PCR protocol listed above with primers pr85 and pr87. Linear DNA templates used for CFE consisted of the gene of interest, codon optimized for *E. coli* expression, flanked by ∼300 bp of upstream and downstream DNA from the pJL1-sfGFP plasmid including the T7 promoter, strong ribosome binding site, and T7 terminator. For the first round of EphB1 redesign, genes encoding the protein of interest were purchased as gene fragments from Twist Biosciences and amplified by PCR as described above. For the second round of EphB1 redesign, genes were amplified from cell cultures as described in “High-throughput data collection.” Reactions expressing EphB1 or redesigned EphB1 variants generally used cell lysates enriched with protein folding chaperones, as described previously.^58^ Reactions were incubated at 30 °C overnight for expression.

#### FluoroTect

15 µL CFE reactions expressing EphB1 or an EphB1 variant were supplemented with 2% v/v FluoroTect reagent (ProMega). After expression, 1.5 µL of RNAse Cocktail (ThermoFisher) was added to CFE reactions and incubated at 37 °C for 10 minutes. 2 µL of the reaction was removed as the total fraction after resuspending and mixing well. CFE reactions were then spun at 18k x g for 15 min at 4 °C. 2 µL of the soluble fraction was removed. Samples were prepared for SDS-PAGE by adding 3.75 µL of 4x Lamelli buffer, 1.5 µL of 1 M DTT, and 7.75 µL of water to the CFE fractions. Samples were denatured at 70 °C for 3 min. Samples were loaded onto a 4-20% tris-glycine gel (Bio-Rad) and ran at 180V for 45 min in 1X tris-glycine running buffer. Gels were imaged using a Typhoon FLA 9000 Imager.

#### PhosphoSens

PhosphoSens (AssayQuant) assays were set up as recommended by the manufacturer in a final volume of 15 µL. All PhosphoSens reactions were run in white, low-volume, 384-well plates (Corning 3824). First, a master reaction mixture consisting of 50 mM HEPES pH 7.5, 0.01% Brij-35, 10 mM MgCl2, 1 mM ATP, 1 mM DTT, 0.55 mM EGTA, and 10µM peptide substrate (AQT0661) was made. 1.27 mL of 10X PhosphoSens buffer (0.5 M HEPES pH 7.5, 0.1% Brij-35, 100 mM MgCl_2_), 1.27 mL of 5.5 mM EGTA, 1.27 mL of 10 mM DTT, 1.27 mL of 10 mM ATP, 1.27 mL of 0.1 mM peptide, and 5.07 mL of water was sufficient master mix for two 384-well plates of PhosphoSens reactions. 13.5 µL of master mix was added to each well in a 384-well plate. 36-fold diluted CFE reactions expressing an EphB1 variant was added to initiate the reaction. Fluorescence values were measured in a 30 °C preheated BioTek Synergy H1 plate reader using an excitation wavelength of 360 nm and an emission wavelength of 492 nm.

The PhosphoSens assay was modified for Michaelis-Menten analysis. 5 mM ATP, corresponding to 10-fold excess of the K_M_ of ATP, was used to ensure that ATP was not limiting.^59^ Peptide stock was made at 5 mM in 20 mM ammonium bicarbonate solution. Purified enzymes were added to PhophoSens assays at a final concentration of 50 nM. Reaction mixtures, prior to adding enzymes, were pre-incubated at 30 °C for 5 min. Enzymes were added, mixed quickly, and immediately measured in a 30 °C preheated plate reader to measure initial rates.

#### Library preparation

Multiplexed gene fragments encoding the redesigned EphB1 variant and 25 bp of sequence homologous to pJL1-HiBiT-EphB1 on the 5’ and 3’ ends were purchased from Twist Biosciences. A linearized backbone of pJL1-HiBiT-EphB1 was prepared by PCR using pr89 and pr90. Successful amplification was confirmed using agarose gel electrophoresis as described previously. The linearized backbone was purified by Qiagen PCR Cleanup and digested using Dpn1 in 1X CutSmart Buffer (NEB) at 37 °C overnight. The redesigned EphB1 fragments were assembled into pJL1 using Gibson Assembly. The library was co-transformed into NEB 5α along with pCDF-PTP1b following manufacturer’s instructions. Serial dilutions of the recovery culture were plated onto LB-Kan/Spec agar plates to confirm > 10X library coverage. The remainder of the recovery culture was inoculated into 10 mL LB-Kan/Spec overnight at 30 °C with 220 rpm shaking. DNA from the culture was miniprepped using Qiagen Miniprep kits and was stored at - 20 °C as the final library.

#### Next generation sequencing

NGS was used to assess the quality of EphB1 libraries. PCR using pr101 and pr104 on pJL1-HiBiT-EphB1 was used to attach NexTera adapters to a 480 bp fragment encoding the redesigned EphB1 fragment. Agarose gel electrophoresis was done to confirm amplification of a single band. A second PCR, using 10% v/v unpurified products from the first PCR as template DNA, was used to attach unique 8-bp indexes to the amplicon using pr107 and pr133. Libraries were cleaned using NucleoMag NGS Cleanup and Size Select Beads (Takara), quantified by Nanodrop, and normalized to ∼ 20 ng/µL. NGS was performed at CZ Biohub on a MiSeq V2 using paired-end 2 x 250 bp runs. Demultiplexed NGS data was processed using CutAdapt.^60^ Sequences were quality-filtered using in-house python scripts by filtering for sequences that were full-length, had a summed error < 0.5, and had > 100 counts. Sequences passing quality-control filters were translated *in silico* and pairwise aligned to a reference database containing all 2000 redesigned variants. Wells containing sequences without any mutations were kept.

#### Split luciferase (HiBiT) quantification

A previously published protocol was adapted for quantification using HiBiT.^37^ The Nano-Glo HiBiT Extracellular Detection System (ProMega) was used to quantify protein expression in CFE reactions. CFE reactions expressing a HiBiT-tagged EphB1 variant were diluted 1000-fold by serial dilution of 1 µL CFE into 35 µL of TBS (50 mM Tris pH 7.5, 300 mM NaCl) and mixed well, and then 1 µL of 36-fold diluted CFE into 27.8 µL of TBS. HiBiT reagent was prepared by mixing 20.2 µL of LgBiT, 40.3 µL of HiBiT substrate, and 1.955 mL HiBiT buffer, which was sufficient for a 384-well plate of HiBiT reactions. 5 µL of HiBit reagent was mixed with 17.5 µL of TBS and 2.5 µL of 1000-fold diluted CFE reactions on ice and immediately placed into a Bio-Tek Synergy H1 plate reader for luminescence measurements. The maximum luminescence was used for all quantification calculations. A standard curve was made using the HiBiT control protein (ProMega) to calculate protein concentration from luminescence.

#### High-throughput data collection

The pooled pJL1-HiBiT-EphB1 variant plasmid library was co-transformed into NEB 5α competent cells along with pCDF-PTP1b such that there were ∼150 colonies per plate. Agar plates were incubated at 30 °C for ∼48 hours to ensure colonies were large enough to be pickable. A QPix-XE automated colony picker was used to pick 1,536 colonies into 16 x 96 deep-well plates containing 1.2 mL of LB-Kan/Spec per well. All plates were incubated at 30 °C with 220 rpm shaking for 24 hours to ensure sufficient growth. 20 µL of saturated cell-culture from four 96-well plates was arrayed into a 384-well plate using a Biomek i7. This was repeated for all sixteen 96-well plates. Cell pellets for long-term storage were made with the remainder of the cultures in 96-well plates by centrifuging the plates at 3k x g for 15 min at 4 °C, decanting the supernatant, and storing at -20 °C. The Biomek i5 equipped with a 384-well head was used for all subsequent plate-to-plate liquid transfers. Linear DNA templates were prepared by first aliquoting 9 µL of PCR premix (containing 2x Master mix, pr85, pr87, and water) into a 384-well plate. 1 µL from saturated cell cultures were added to 384-well PCR plates and were thermocycled as listed above. Random wells were spot-checked by agarose gel electrophoresis to confirm amplification. 6 µL of CFE reaction mixture was then aliquoted into another 384-well plate, and 1 µL of PCR product was added to CFE reactions and well-mixed to express EphB1 variants. CFE reactions were incubated overnight at 30 °C in a shaking incubator. 1 µL from the CFE reactions was diluted into 35 µL of TBS for PhosphoSens assays, and the 36-fold diluted CFE reactions were serially diluted into 27.8 µL of TBS to make a 1000-fold diluted CFE reaction suitable for HiBiT quantification. 1.5 µL of the 36-fold diluted CFE reaction was mixed with 13.5 µL of PhosphoSens master mix and mixed by pipetting to initiate the reaction. 2.5 µL of the 1000-fold diluted CFE reaction was mixed with 22.5 µL of HiBiT master mix and mixed by pipetting for split luciferase complementation.

All four 384-well plates were barcoded to identify each variant in each well. A 480-bp amplicon encoding the redesigned fragment was lifted from each variant using PCR with pr101 and pr104, which also attached NexTera adapters. Random spot checks of wells confirmed successful amplification. Unique 12-bp dual indexes were attached to each amplicon using PCR with indexing primers generously provided by CZ Biohub. Random spot checks confirmed amplification. All 1,536 wells were pooled, cleaned up using NucleoMag NGS Cleanup and Size Select Beads (Takara), quantified by Nanodrop, and normalized to ∼ 20 ng/µL. 15M reads were collected on a MiSeq V2 on a paired 2 x 250 bp run. Data was analyzed as described above.

#### Testing model predictions

Unseen variants were ordered as gene fragments from Twist Biosciences with overhangs appropriate for Gibson assembly. Linearized backbone was prepared using PCR on pJL1-HiBiT-EphB1 with pr89 and pr90. Gibson assembly was done as previously stated to obtain sequence-verified plasmids with redesigned EphB1 variants. A linear DNA template was amplified using PCR with pr85 and pr87. EphB1 variants were expressed in CFE as previously stated. PhosphoSens and HiBiT quantification was performed as previously stated.

#### Protein expression and purification

*E. coli* BL21 (DE3) was co-transformed with pET-EphB1 and with pCDF-PTP1b using manufacturer protocols. Cells were plated onto LB agar with appropriate antibiotics and incubated at 30 °C overnight. Single colonies were inoculated into 5 mL of LB with appropriate antibiotics and incubated at 30 °C overnight with 220 rpm shaking. For data in **Fig S2**, 5 mL of TB with appropriate antibiotics were inoculated with a 1:100 dilution of the overnight culture and grown at 30 °C. At OD_600_ = 0.6 – 0.8 or after 8 hours of growth (as cells expressing EphB1 without PTP1b exhibited a noticeable growth defect), cells were induced with 1 mM IPTG, transferred to an 18 °C shaker, and incubated overnight with 220 rpm shaking. After overnight growth, 2 µL from each cell culture was analyzed by SDS-PAGE for protein expression. For data in **Fig S12**, 50 mL of Overnight Express TB Media (Millipore) with appropriate antibiotics was inoculated with a 1:100 dilution of overnight culture. Cultures were grown at 37 °C for ∼ 4 hours with 220 rpm shaking until cultures grew to OD_600_ = 0.6 – 0.8. Cultures were transferred to an 18 °C pre-chilled incubator at 220 rpm shaking and incubated overnight. After ∼20 hours of growth, cells were pelleted and stored at -20 °C.

Proteins were purified by IMAC gravity purification. Cell pellets were thawed at room temperature and resuspended in 2 mL of HBS (50 mM HEPES pH 7.5, 300 mM NaCl) with 10 mM imidazole pH 7.5 per gram of wet cell mass by vortexing. Cells were lysed by adding 1 mg/mL of lysozyme and 5 U of Benzonase, incubating at room temperature with end-over-end shaking for 30 min at room temperature, and sonicating for 3 min with 5 s ON / 5s OFF cycles at 20% amplitude. Cell lysate was centrifuged at 20k x g for 20 min at 4 °C to remove insoluble components. Supernatant was filtered through a 0.45 µm filter. Soluble lysate was mixed with pre-equilibrated NiNTA resin (Qiagen) and incubated at 4 °C with end-over-end shaking for 1 hr. All proteins were purified using batch purification. Resins were washed 5X by resuspending in 10 mL of wash buffer (HBS + 20 mM imidazole pH 7.5), pelleting the resin by centrifugation at 5k x g for 2 min at 4 °C, and decanting the supernatant. Resins were transferred to a gravity flow column and protein were eluted in 5 mL of elution buffer (HBS + 250 mM imidazole pH 7.5) split into 5 fractions. Fractions were analyzed by SDS-PAGE. 1 µL of protein from each fraction was mixed with 3.75 µL of 4X Lamelli buffer, 1.5 µL of 1 M DTT, and 8.75 µL of water. Samples were denatured at 95 °C for 10 min. Samples were loaded onto a 4-20% tris-glycine gel (Bio-Rad) and ran at 180V for 45 min in 1X tris-glycine running buffer. Gels were stained in InstantBlue (Bulldog Bio) and destained in water overnight. Protein-containing fractions were consolidated and dialyzed against 4L of HBS overnight at 4 °C.

Proteins were further purified using FPLC. Proteins were concentrated to 0.5 mL using Amicon 3.5 kDa ultracentrifugation filters. FPLC was performed using an Akta PURE connected to a Cytiva Superdex Increase 200 10/300 column using HBS as running buffer. Bio-Rad Gel Filtration Standard was used to estimate the elution profile of the column. All proteins eluted at the expected molecular weight. Fractions corresponding to the monomeric peak were collected, analyzed again by SDS-PAGE to confirm purity, and stored at 4 °C for downstream assays. Absorbance at 280 nm was measured by Nanodrop and protein concentrations were calculated using molecular weights and extinction coefficients calculated by ExPasy ProtParam. All proteins were freshly purified and used within a week.

### Computational Methods

#### General methods

All plotting was done in Python either with seaborn or with matplotlib. All residue numbering for EphB1 is consistent with UniProt P54762 unless otherwise stated.

#### EphB1 kinase redesign

Protein kinases were redesigned using the strategy outlined in Sumida *et al*.^31^ First, evolutionary conserved residues in protein tyrosine kinases were identified. Sequences for the catalytic domain of protein tyrosine kinases were retrieved from InterPro using ID IPR020635. 500 sequences, including the EphB1 sequence, were randomly selected and aligned using the MUSCLE webserver (https://www.ebi.ac.uk/jdispatcher/msa/muscle) to make an MSA. For columns where the WT EphB1 was not a gap, the frequency of each amino acid was calculated. If the frequency of any amino acid within the column exceeded the evolutionary conservation thresholds, the corresponding residue in EphB1 was fixed during redesign. This process was repeated four times with other sets of 500 randomly selected sequences. All residues that were below the evolutionary conservation thresholds were set as designable. Next, we identified residues in the active sites of EphB1. We used a representative crystal structure of EphB1 (PDBID: 7kpm) and identified all residues containing a C_ß_ within 7Å of the ADP molecule using PyMOL. To identify positions that may interact with the peptide substrate, we aligned EphB1 with EphA3, a closely related homolog whose structure was solved in complex with a peptide substrate (PDBID: 3fxx). We aligned EphB1 using the align function in PyMOL. Peptide-binding residues were defined as those with a C_ß_ within 7Å of the peptide substrate. As an alternative strategy, we also fixed all residues within the core and boundary of EphB1 and redesigned the surface (see next section).

~~~
7kpm sequence (residues 611 – 889) AKEIDVSFVKIEEVIGAGEFGEVYKGRLKLPGKREIYVAIKTLKAGYSEKQRRDFLSEASIMGQFDHPNIIRLEGVVTKSRPVMIITEFMENGALDSFLRQNDGQFTVIQLVGMLRGIAAGMKYLAEMNYVHRDLAARNILVNSNLVCKVSDFGLSRYLQDDTSDPTYTSSLGGKIPVRWTAPEAIAYRKFTSASDVWSYGIVMWEVMSFGERPYWDMSNQDVINAIEQDYRLPPPMDCPAALHQLMLDCWQKDRNSRPRFAEIVNTLDKMIRNPASLK
~~~

Residues > 50% conservation (1-indexed for pMPNN and F2s inputs):

~~~
[15, 16, 17, 18, 19, 20, 21, 23, 26, 38, 39, 40, 41, 43, 44, 54, 55, 56,
58, 59, 61, 62, 67, 68, 69, 70, 71, 73, 75, 76, 77, 78, 82, 85, 86, 87, 88,
89, 90, 93, 94, 95, 98, 99, 100, 111, 117, 118, 119, 121, 122, 124, 125,
131, 132, 133, 134, 135, 136, 137, 138, 139, 141, 142, 145, 149, 150, 151,
152, 153, 154, 155, 156, 157, 158, 159, 160, 161, 162, 163, 164, 165, 166,
167, 168, 169, 170, 171, 172, 173, 174, 175, 176, 177, 178, 179, 180, 181,
182, 183, 184, 185, 186, 187, 189, 191, 192, 195, 196, 197, 198, 199, 200,
201, 202, 205, 206, 209, 211, 213, 214, 215, 219, 220, 221, 223, 230, 231,
232, 233, 234, 236, 239, 240, 243, 244, 246, 247, 250, 251, 255, 258, 259,
261, 265]
~~~

Residues > 75% conservation (1-indexed for pMPNN and F2s inputs):

~~~
[15, 16, 17, 18, 19, 20, 21, 23, 38, 39, 41, 43, 58, 62, 67, 71, 73, 75,
82, 85, 87, 88, 89, 90, 93, 94, 95, 99, 111, 119, 121, 122, 124, 125, 132,
133, 134, 135, 136, 137, 138, 139, 141, 149, 151, 152, 153, 154, 155, 156,
157, 158, 159, 160, 161, 162, 163, 164, 165, 166, 167, 168, 169, 170, 171,
172, 173, 174, 175, 176, 177, 178, 179, 180, 182, 184, 186, 187, 192, 195,
196, 197, 198, 201, 205, 206, 211, 213, 214, 215, 219, 220, 221, 230, 231,
232, 233, 236, 239, 247, 250, 251, 258, 259, 261]
~~~

Residues > 95 % conservation (1-indexed for pMPNN and F2s inputs):

~~~
[15, 16, 17, 18, 19, 20, 21, 23, 39, 41, 58, 71, 85, 87, 88, 89, 90, 93,
94, 95, 125, 132, 133, 134, 136, 138, 139, 141, 151, 152, 153, 154, 155,
156, 157, 158, 159, 160, 161, 162, 163, 164, 165, 166, 167, 168, 169, 170,
171, 172, 173, 174, 175, 176, 177, 178, 179, 180, 184, 186, 187, 195, 196,
198, 205, 206, 213, 214, 219, 220, 221, 236, 250, 258, 259, 261]
~~~

Core and boundary (1-indexed for pMPNN and F2s inputs):

~~~
[9, 15, 16, 18, 19, 20, 21, 23, 24, 26, 28, 38, 39, 40, 41, 43, 47, 51, 54
, 55, 58, 59, 61, 62, 65, 67, 69, 70, 71, 73, 74, 75, 76, 77, 82, 83, 84,
85, 86, 87, 89, 90, 92, 93, 94, 95, 96, 97, 98, 99, 106, 110, 111, 114, 11
5, 117, 118, 119, 121, 122, 123, 124, 125, 126, 128, 130, 131, 132, 133, 1
34, 135, 136, 137, 139, 140, 141, 142, 146, 148, 149, 150, 151, 152, 153,
154, 155, 156, 158, 159, 163, 168, 169, 171, 174, 175, 176, 177, 178, 179,
180, 181, 182, 183, 184, 185, 186, 191, 193, 194, 195, 196, 197, 198, 199,
200, 201, 202, 203, 204, 205, 206, 207, 208, 209, 210, 211, 212, 214, 215,
218, 220, 222, 223, 226, 227, 231, 232, 233, 236, 238, 239, 243, 244, 246,
247, 250, 251, 252, 255, 258, 259, 261, 264, 265, 267]
~~~

EphB1 variants were redesigned using ProteinMPNN (https://github.com/dauparas/ProteinMPNN)^19^ and Frame2seq (https://github.com/dakpinaroglu/Frame2seq).^20^ The 7kpm structure was used as the input for all redesign simulations. Residues 602-610 and 890-896 were manually added back to redesigned sequences as they are not included in the 7kpm structure. In the pilot study, the full gene was redesigned using pMPNN or soluble pMPNN. 300 sequences, equally split between temperatures = 0.1, 0.2, and 0.3, were made for every set of redesignable residues (< 50%, < 75%, < 95%, and surface), resulting in a total of 2,400 sequence designs. In the second-generation library, each round of design redesigned all residues < 50% conservation and a random subset of residues between 50 – 75% conservation for redesign. 300 sequences were made using pMPNN (temperatures = 0.1, 0.3, and 0.5, 100 sequences each) and with Frame2seq (temperatures = 0.3, 0.65, 1, 100 sequences each). This process was repeated 100x for a total of 60,000 sequence designs. pMPNN and Frame2seq do not design unmodeled residues (they are masked as ‘X’ during redesign), so these residues were manually reverted to the native amino acid in EphB1.

Sequences were scored using ESM-1v and Frame2seq (only in the pilot study). ESM-1v scores were calculated using the protein_gibbs_sampler package (https://github.com/seanrjohnson/protein_gibbs_sampler). The ‘likelihood_esm.py’ script was used to calculate ESM-1v scores for all sequences using the --masking off flag, as described in previous works.^33^ Frame2seq scores were calculated using the score function of Frame2seq as outlined in the github repository.

#### AlphaFold2 structure prediction

We used LocalColabFold (https://github.com/YoshitakaMo/localcolabfold) to predict structures for kinase sequences with MSA, no templates, all 5 models, and 3 recycles. The structure of the EphB1 kinase was not predicted well without MSA (Cα RMSD: 16.8 Å, pLDDT = 38.8 from the best model). We used single-sequence mode (no MSA) to screen redesigned kinases in the pilot study to assess if the structures of any redesigned variants would be predicted more confidently. In the first round of redesign, we selected the best predicted sequence that scored moderately with a pLDDT of ∼80 and Cα RMSD of 1.93 Å from the best model without MSA.

#### EphB1 structural analysis

We used the LayerSelector within PyRosetta to identify the core, boundary, and surface residues of EphB1 (https://www.pyrosetta.org/home). As some residues were unmodelled in the experimental crystal structure of EphB1 (7kpm), we used an AlphaFold2 prediction for the structure. The prediction has < 1 Å Cα RMSD to the experimental structure over the well-resolved regions. The predicted structure was relaxed using PyRosetta FastRelax using default settings (ref2015 score function, 5 repeats) using the following code block below. The LayerSelector was applied to the relaxed structure to identify residues within each layer of the structure.

~~~
pose = pyrosetta.io.pose_from_pdb(’ephb1_af2.pdb’)
sfxn = get_score_function()
fr = pyrosetta.rosetta.protocols.relax.FastRelax()
fr.set_scorefxn(sfxn)
fr.apply(pose)
~~~

To identify key regulatory features of protein kinases such as the R spine and C spine, we aligned a structure of Protein Kinase A (PDBID: 4wb5) with the relaxed EphB1 structure described above and used the numbering provided in ref.^61^ to identify R spine and C spine residues. DSSP was used to identify residues in the αC helix using pyrosetta.rosetta.protocols.mover.DsspMover. The activation loop was defined as all residues between (and including) the DFG motif and the APE motif. Distances from structural features were calculated by measuring the distance Cα distance between all residues in each feature with the Cα of the residue of interest and recording the shortest distance.

#### EphB1 sequence analysis

Pairwise mutations were calculated using an in-house python script. The theoretical complete size of the sequence space explored by the redesigned kinases was calculated by multiplying the number of mutations at each redesigned site. The total number of unique mutations was calculated by adding the number of unique mutations at each redesigned site.

Principal component analysis of OHE protein sequences or ESM2 embeddings were performed using sklearn. Sequences were one-hot encoded using an in-house python script. ESM2 embeddings and ESM-1v embeddings (both from the 650 M parameter models) were computed using the esm-extract script with redesigned EphB1 variants inputted in fasta format (https://github.com/facebookresearch/esm/tree/main), using the mean of embeddings from layers 0, 32, and 33 as described in the github repository. PCA was performed on OHE sequences and on ESM embeddings using default settings.

UMAP analysis was done using the umap-learn package (https://github.com/lmcinnes/umap). All sequences of natural protein tyrosine kinase domains from InterPro were first clustered with the MMseqs2 package (https://github.com/soedinglab/mmseqs2) using the easy-cluster function with default settings. This resulted in 314 representative sequences. ESM2 embeddings for representative kinases were computed as described above. UMAP was performed on this dataset, along with with active redesigned EphB1 variants identified in the experimental analysis (defined as those with specific activities greater than 15,000 RFU min^-1^ µM^-1)^. UMAP was done with default settings except that n_neighbors = 8.

Categorical entropy at designed position was calculated using physiochemical categories of each mutation (Hydrophobic: A,L,I,V,M; Aromatic: F,W,Y; Polar: S,T,N,Q; Charged: D,E,K,R; Glycine: G, Proline: P, Cysteine: C, Histidine: H).

#### Michaelis-Menten fitting

Initial rates were calculated by fitting a linear regression line on the first 90 seconds of the enzyme progress curve for each substrate concentration using scipy.stats.linregress. The qualities of the fits were assessed by calculating pearson r values. Data were fitted to the Michaelis-Menten equation using scipy.optimize.curve_fit with initial guesses of k_cat_ = 0.6 s^-1^ and K_M_ = 40 µM.

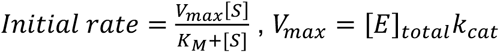

One standard deviation of the k_cat_ and K_M_ parameters was calculated by taking the square root of the diagonals of the estimated approximate covariance of the parameters (https://docs.scipy.org/doc/scipy/reference/generated/scipy.optimize.curve_fit.html).

#### Data processing

Raw activity rates were normalized by the corresponding measured concentration to calculate the specific activity for each design. Negative specific activity values, which were small in magnitude and likely reflective of assay noise, were clipped to zero. In cases where a design had replicate measurements, the specific activity values were averaged to a single rate per design. As the distribution of specific activities was right-skewed, a Yeo-Johnson transformation was applied prior to model fitting.

#### Feature engineering

Multiple feature representations were evaluated to capture sequence and/or structural properties of each design. Amino acid sequences in the redesignable region were first encoded using one-hot encoding. In addition, we also encoded the sequences using Georgiev encodings to include physicochemical characteristics.^40^ ESM-1v embeddings were also calculated as described above to incorporate implicit structural and evolutionary information.

These sequence-based features were then combined with predictions of fitness. We trained an EVmutation pairwise epistatic model using the EVcouplings webserver (https://evcouplings.org) using the input sequence of the tyrosine kinase domain for wild-type EphB1 with a bitscore inclusion threshold of 0.7. This model was used to calculate the change in statistical energy compared to the wild-type EphB1 as a feature representing evolutionary fitness.

We also input the structure used during design (PDB ID: 7kpm) into ThermoMPNN, a deep learning method trained to predict the change in thermostability caused by point mutations to a given sequence.^3^ The resulting mutational ddG matrix was used to estimate the ddG value compared to wild-type EphB1 for each design to incorporate structure-based stability information as well.

#### Machine learning

Sequence featurization was done by concatenating one or more fitness predictions (EVcouplings score, ThermoMPNN ddG) with one sequence-level amino acid representation (one-hot, Georgiev, or ESM-1v embeddings). Continuous features (ESM-1v embeddings, EVCouplings score, ThermoMPNN ddG) were first standardized. The L1 and L2 regularization strengths for lasso and ridge regression, respectively, were determined via hyperparameter tuning within each fold during cross-validation. Random Forest regressor model was implemented using the default settings from scikit-learn except with 50 estimators. We assessed model performance using 5-fold cross validation and report the average Spearman r and MSE across all splits.

A ridge regression model using one-hot encoded sequences was trained to facilitate interpretation of the effect of individual mutations on specific activity. To evaluate the relative importance of each position in predicting activity, the absolute values of the model coefficients were fitted, and their sum across all amino acid possibilities at that position was taken as a proxy for positional importance, i.e. large coefficient sums indicated that the amino acid identity at that position was informative to the model. To analyze the dependency of model performance on the number of positions included, individual ridge regression models were trained on progressively larger subsets of residues. Starting with the single most informative position, one additional position was added at each iteration in descending order of their positional importance. Model performance for each subset was evaluated using 5-fold cross validation as described previously.

The ridge regression model trained on all labeled one-hot encoded sequences was used to predict the activity of unseen variants. The redesigned regions of all unseen variants in the library were one-hot encoded. Variants were excluded if they contained any individual mutation that was not present in the training data (n = 235).

#### Potts models

Potts models were fit using scripts from https://github.com/sokrypton/GREMLIN_CPP/blob/master/GREMLIN_TF.ipynb.^46,62^ This implementation uses pseudolikelihood optimization to fit the one- and two-body terms of the Potts model. The Potts model was trained on an MSA of Eph receptor tyrosine kinase homologues. The MSA was obtained using localcolabfold (https://github.com/YoshitakaMo/localcolabfold) with the WT EphB1 sequence as input, which uses MMseqs2 to search Uniref30, BFD, and MGnify to find sequence homologues.^63^ The MSA was converted from an a3m file to a fasta file using https://bioi2.i2bc.paris-saclay.fr/msa-tools/format-conversion/.

Natural Eph receptor tyrosine kinase homologues used to compare Potts model performance against redesigned variant Potts models were retrieved from InterPro using the ID IPR050449. Sequences annotated as “Ephrin receptor tyrosine kinases” or “Eph receptor tyrosine kinases” were kept. Sequences were aligned to the MSA used to train the Potts model using MAFFT (https://mafft.cbrc.jp/alignment/software/) with flags:

~~~
mafft --inputorder --keeplength --compactmapout --add new_sequences --auto input
~~~

537 sequences, matching the number of redesigned sequences, were randomly selected and scored using the trained Potts model. Sequences that contained > 10% gaps, contained unknown residues (“X”), or were duplicated within the MSA used during Potts model training were removed. This was repeated three times to confirm that the results were robust.

#### Weblogos

MSAs were created with MUSCLE5 using sequences retrieved as previously described (representative sequences described under UMAP, and Eph tyrosine kinase homologues described under Potts models). Columns in the MSA with > 75% gaps were removed. Weblogos were created using Logomaker (https://github.com/jbkinney/logomaker).

## Supporting information

DNA sequences

Raw uncropped gels

## Acknowledgements

We thank A. Li and D. Muir for helpful discussions on computational modeling, S. Ghaffari-Kashani for insight into kinase assays, S. Crilly for help with NGS, E. Zhu and I. Alfonso for laboratory management, members of the Kortemme Lab for helpful discussions, and Prof. M. Jewett and his lab for providing initial CFE reagents as benchmarks. We thank J. Posner and the Pinney lab for training and use of the QPix-XE colony picker. This work made use of the UCSF Wynton HPC, which is supported by UCSF Faculty and UCSF institutional funds, and we are thankful to the UCSF Wynton team for technical support. We gratefully acknowledge CZ Biohub for NGS support and for generously providing UDI primers. This work made use of the Biomek i5, Biomek i7, and 384-well head thermocyclers housed at the UCSF CAT, which is supported by UCSF PBBR, RRP IMIA, and NIH 1S10OD028511-01 grants.

## Author Contributions

K.S. conceived the idea for the project. K.S., A.B.G., and T.K. developed the conceptual approach. K.S. performed all design simulations with contributions from D.A., developed the CFE workflow, performed all biochemical experiments, and analyzed the data. A.B.G. conceptualized and performed model training, and analyzed the data. T.K. provided guidance and resources. K.S., A.B.G., and T.K. wrote the manuscript. All authors read and commented on the manuscript. Conceptualization: K.S., T.K., Methodology: K.S., A.B.G., D.A., Software: K.S., Validation: K.S., Formal analysis: K.S., A.B.G., Investigation: K.S., Resources: K.S., Data curation: K.S., Writing: K.S., A.B.G., T.K., Visualization: K.S., Supervision: T.K., Project administration: T.K., Funding acquisition: T.K..

## Data and materials availability

All code will be made available on a github repository. Data are available upon request.

