## Supplementary Figures and Tables for "A combinatorial mutational map of active non-native protein kinases by deep learning guided sequence design"

**Table S1**: Redesigned EphB1 variants from pMPNN pilot library. Variants are named according to an A-B-C-D scheme, where A refers to the fixed residue set (conservation threshold, surface design) and the version of pMPNN used (soluble or default), B refers to the PDB input, C refers to the design temperature used in pMPNN, and D is a sample number.

| ID | Selection Criterion | Active? | Mutations from WT |
|---|---|---|---|
| 95-7kpm-0.3-53 | AF2 pLDDT | N | 114 |
| 50-7kpm-0.3-2 | max esm-1v | Y | 87 |
| 75-7kpm-0.1-7 | max esm-1v | Y | 100 |
| 95-7kpm-0.2-55 | max esm-1v | N | 112 |
| sol50-7kpm-0.3-89 | max esm-1v | Y | 87 |
| sol75-7kpm-0.2-14 | max esm-1v | N | 100 |
| sol95-7kpm-0.1-82 | max esm-1v | N | 102 |
| surface_sol-7kpm-0.1-92 | max esm-1v | N | 72 |
| surface-7kpm-0.3-22 | max esm-1v | N | 71 |
| 50-7kpm-0.2-89 | max f2s | Y | 86 |
| 75-7kpm-0.2-18 | max f2s | N | 91 |
| 95-7kpm-0.1-74 | max f2s | N | 105 |
| sol50-7kpm-0.2-70 | max f2s | N | 86 |
| sol75-7kpm-0.2-57 | max f2s | Y | 101 |
| sol95-7kpm-0.3-48 | max f2s | N | 94 |
| surface_sol-7kpm-0.3-96 | max f2s | N | 69 |
| surface-7kpm-0.1-74 | max f2s | N | 68 |
| 50-7kpm-0.3-66 | min esm-1v | N | 82 |
| 75-7kpm-0.3-70 | min esm-1v | N | 104 |
| 95-7kpm-0.3-3 | min esm-1v | N | 113 |
| sol50-7kpm-0.2-71 | min esm-1v | N | 89 |
| sol75-7kpm-0.1-19 | min esm-1v | N | 101 |
| sol95-7kpm-0.3-26 | min esm-1v | N | 110 |
| surface_sol-7kpm-0.3-13 | min esm-1v | N | 73 |
| surface-7kpm-0.3-92 | min esm-1v | N | 73 |

**Table S2:** Counts of all residues at all redesigned positions in the redesigned EphB1 variants tested in our workflow. Amino acids are listed in one-letter code.

| Position | Amino Acid | Counts |
|---|---|---|
| 661Q | D | 28 |
| | E | 371 |
| | S | 1 |
| | Q | 135 |
| | A | 2 |
| 662R | G | 1 |
| | E | 45 |
| | K | 373 |
| | R | 65 |
| | S | 2 |
| | Q | 31 |
| | A | 6 |
| | L | 4 |
| | I | 10 |
| 663R | E | 234 |
| | K | 166 |
| | R | 18 |
| | S | 4 |
| | T | 5 |
| | N | 6 |
| | Q | 13 |
| | A | 80 |
| | L | 4 |
| | I | 2 |
| | V | 5 |
| 664D | D | 313 |
| | E | 124 |
| | K | 13 |
| | R | 1 |
| | S | 2 |
| | Q | 6 |
| | A | 76 |
| | L | 2 |
| 666L | L | 535 |
| | I | 1 |

| | | |
|---|---|---|
| | M | 1 |
| 667S | G | 1 |
| | H | 4 |
| | D | 10 |
| | E | 67 |
| | K | 191 |
| | R | 16 |
| | S | 12 |
| | N | 140 |
| | Q | 71 |
| | A | 22 |
| | L | 1 |
| | M | 1 |
| | Y | 1 |
| 670S | H | 4 |
| | D | 11 |
| | E | 198 |
| | K | 61 |
| | R | 45 |
| | S | 20 |
| | T | 170 |
| | N | 2 |
| | Q | 6 |
| | A | 8 |
| | L | 3 |
| | I | 1 |
| | V | 3 |
| | M | 1 |
| | F | 1 |
| | Y | 3 |
| 671I | T | 5 |
| | L | 66 |
| | I | 433 |
| | V | 32 |
| | M | 1 |
| 673G | G | 178 |
| | S | 238 |
| | N | 4 |
| | A | 117 |

| | | |
|---|---|---|
| 674Q | E | 12 |
| | K | 294 |
| | R | 35 |
| | T | 1 |
| | N | 1 |
| | Q | 191 |
| | L | 1 |
| | M | 2 |
| 676D | D | 287 |
| | S | 2 |
| | N | 246 |
| | Q | 2 |
| 678P | P | 531 |
| | D | 3 |
| | E | 2 |
| | N | 1 |
| 679N | H | 5 |
| | N | 532 |
| 680I | I | 535 |
| | V | 2 |
| 682R | K | 500 |
| | R | 20 |
| | T | 3 |
| | N | 1 |
| | Q | 12 |
| | A | 1 |
| 684E | H | 5 |
| | E | 5 |
| | K | 6 |
| | R | 3 |
| | Q | 1 |
| | L | 421 |
| | I | 21 |
| | V | 2 |
| | F | 5 |
| | W | 1 |
| | Y | 67 |
| 686V | I | 3 |
| | V | 534 |

| | | |
|---|---|---|
| | I | 45 |
| | V | 492 |
| 688T | T | 532 |
| | I | 3 |
| | V | 2 |
| 689K | H | 5 |
| | D | 5 |
| | E | 8 |
| | K | 379 |
| | R | 13 |
| | S | 52 |
| | T | 54 |
| | N | 10 |
| | Q | 7 |
| | A | 1 |
| | L | 2 |
| | V | 1 |
| 690S | H | 5 |
| | D | 6 |
| | E | 255 |
| | K | 12 |
| | R | 4 |
| | S | 26 |
| | T | 208 |
| | Q | 16 |
| | A | 5 |
| 691R | H | 6 |
| | D | 21 |
| | E | 381 |
| | K | 30 |
| | R | 12 |
| | S | 32 |
| | T | 4 |
| | N | 10 |
| | Q | 27 |
| | A | 3 |
| | L | 5 |
| | F | 2 |
| | Y | 4 |

| | | |
|---|---|---|
| 693V | P | 2 |
| | H | 1 |
| | K | 1 |
| | Q | 2 |
| | A | 1 |
| | L | 249 |
| | I | 46 |
| | V | 232 |
| | M | 2 |
| | F | 1 |
| 694M | H | 1 |
| | K | 5 |
| | R | 5 |
| | L | 15 |
| | M | 506 |
| | Y | 5 |
| 696I | I | 514 |
| | V | 23 |
| 701E | P | 8 |
| | D | 5 |
| | E | 472 |
| | K | 13 |
| | S | 12 |
| | Q | 4 |
| | A | 23 |
| 702N | H | 68 |
| | K | 215 |
| | R | 33 |
| | N | 193 |
| | Q | 9 |
| | L | 14 |
| | Y | 5 |
| 706D | G | 29 |
| | H | 27 |
| | D | 37 |
| | E | 2 |
| | K | 89 |
| | R | 114 |
| | S | 3 |

| | | |
|---|---|---|
| | T | 2 |
| | N | 6 |
| | Q | 24 |
| | A | 1 |
| | L | 182 |
| | F | 1 |
| | Y | 20 |
| 707S | G | 15 |
| | P | 3 |
| | H | 3 |
| | D | 270 |
| | E | 45 |
| | K | 10 |
| | R | 11 |
| | S | 99 |
| | T | 36 |
| | N | 30 |
| | Q | 9 |
| | A | 6 |
| 708F | F | 445 |
| | Y | 92 |
| 710R | K | 5 |
| | R | 521 |
| | Q | 11 |
| 711Q | G | 6 |
| | D | 13 |
| | E | 124 |
| | K | 30 |
| | R | 25 |
| | S | 139 |
| | N | 106 |
| | Q | 39 |
| | A | 55 |
| 712N | H | 79 |
| | N | 458 |
| 713D | E | 2 |
| | K | 102 |
| | R | 233 |
| | T | 2 |

| | | |
|---|---|---|
| | Q | 4 |
| | I | 32 |
| | V | 161 |
| | Y | 1 |
| 714G | G | 535 |
| | R | 2 |
| 715Q | G | 5 |
| | H | 2 |
| | D | 1 |
| | E | 42 |
| | K | 360 |
| | R | 16 |
| | S | 19 |
| | Q | 87 |
| | A | 5 |
| 716F | L | 243 |
| | F | 252 |
| | Y | 42 |
| 717T | G | 1 |
| | S | 423 |
| | T | 113 |
| 718V | T | 84 |
| | I | 272 |
| | V | 181 |
| 719I | P | 1 |
| | E | 7 |
| | K | 4 |
| | R | 4 |
| | T | 5 |
| | N | 1 |
| | Q | 9 |
| | L | 403 |
| | I | 13 |
| | V | 3 |
| | M | 82 |
| | Y | 5 |
| 720Q | D | 2 |
| | E | 268 |
| | R | 1 |

| | | |
|---|---|---|
| | T | 4 |
| | Q | 261 |
| | Y | 1 |
| 722V | T | 1 |
| | L | 11 |
| | I | 7 |
| | V | 518 |
| 723G | G | 13 |
| | H | 12 |
| | E | 18 |
| | K | 110 |
| | R | 267 |
| | S | 10 |
| | N | 80 |
| | Q | 23 |
| | A | 1 |
| | L | 2 |
| | M | 1 |
| 724M | I | 165 |
| | M | 372 |
| 725L | A | 1 |
| | L | 529 |
| | M | 7 |
| 726R | H | 1 |
| | K | 23 |
| | R | 176 |
| | Q | 2 |
| | L | 335 |
| 727G | G | 385 |
| | D | 125 |
| | E | 2 |
| | S | 9 |
| | N | 15 |
| | Q | 1 |
| 728I | A | 1 |
| | I | 536 |
| 730A | D | 4 |
| | E | 66 |
| | K | 197 |

| | | |
|---|---|---|
| | R | 224 |
| | S | 9 |
| | N | 1 |
| | Q | 35 |
| | L | 1 |
| 733K | C | 1 |
| | H | 1 |
| | D | 75 |
| | E | 232 |
| | K | 28 |
| | R | 9 |
| | S | 3 |
| | T | 1 |
| | N | 99 |
| | Q | 39 |
| | A | 42 |
| | L | 5 |
| | M | 2 |
| 736A | C | 14 |
| | H | 49 |
| | D | 1 |
| | S | 10 |
| | N | 1 |
| | A | 462 |
| 737E | G | 1 |
| | D | 24 |
| | E | 173 |
| | K | 83 |
| | R | 8 |
| | S | 179 |
| | N | 6 |
| | Q | 51 |
| | A | 9 |
| | L | 3 |
| 738M | G | 1 |
| | H | 59 |
| | E | 1 |
| | K | 270 |
| | R | 15 |

| | | |
|---|---|---|
| | N | 171 |
| | Q | 19 |
| | M | 1 |
| 739N | G | 88 |
| | H | 10 |
| | D | 21 |
| | E | 10 |
| | K | 150 |
| | R | 12 |
| | S | 1 |
| | N | 227 |
| | Q | 16 |
| | A | 2 |
| 740Y | L | 1 |
| | I | 1 |
| | V | 2 |
| | F | 210 |
| | Y | 323 |
| 741V | I | 2 |
| | V | 535 |
| 750I | L | 11 |
| | I | 517 |
| | V | 9 |
| 752V | L | 1 |
| | I | 83 |
| | V | 453 |
| 753N | D | 528 |
| | N | 9 |
| 754S | G | 3 |
| | H | 1 |
| | D | 12 |
| | E | 297 |
| | K | 63 |
| | R | 1 |
| | S | 52 |
| | T | 1 |
| | N | 1 |
| | Q | 7 |
| | A | 96 |

| | | |
|---|---|---|
| | I | 1 |
| | V | 2 |
| 755N | D | 31 |
| | E | 11 |
| | K | 6 |
| | S | 2 |
| | T | 1 |
| | N | 483 |
| | Q | 3 |
| 756L | D | 1 |
| | K | 1 |
| | N | 2 |
| | L | 522 |
| | I | 1 |
| | M | 9 |
| | Y | 1 |
| 757V | C | 1 |
| | H | 1 |
| | E | 3 |
| | K | 16 |
| | R | 22 |
| | T | 161 |
| | N | 279 |
| | Q | 38 |
| | I | 5 |
| | V | 10 |
| | M | 1 |
| 758C | C | 290 |
| | T | 2 |
| | A | 236 |
| | V | 9 |
| 760V | I | 161 |
| | V | 376 |
| 791T | S | 10 |
| | T | 464 |
| | L | 41 |
| | M | 22 |
| 793P | P | 527 |
| | E | 7 |

| | | |
|---|---|---|
| | L | 3 |
| 795A | A | 514 |
| | L | 15 |
| | M | 3 |
| | F | 5 |
| 798Y | G | 257 |
| | H | 19 |
| | E | 23 |
| | K | 4 |
| | R | 3 |
| | S | 1 |
| | T | 1 |
| | N | 213 |
| | Q | 13 |
| | A | 1 |
| | Y | 2 |
| 799R | G | 151 |
| | H | 1 |
| | E | 4 |
| | K | 15 |
| | R | 316 |
| | S | 3 |
| | N | 12 |
| | Q | 5 |
| | A | 29 |
| | L | 1 |
| 800K | H | 1 |
| | D | 187 |
| | E | 36 |
| | K | 20 |
| | R | 11 |
| | S | 3 |
| | T | 198 |
| | N | 42 |
| | Q | 13 |
| | A | 1 |
| | L | 5 |
| | I | 16 |
| | V | 4 |

| | | |
|---|---|---|
| 801F | P | 9 |
| | H | 1 |
| | F | 435 |
| | W | 33 |
| | Y | 59 |
| 803S | G | 131 |
| | P | 228 |
| | C | 16 |
| | H | 7 |
| | D | 6 |
| | E | 5 |
| | K | 17 |
| | R | 21 |
| | S | 17 |
| | T | 6 |
| | N | 1 |
| | Q | 4 |
| | A | 53 |
| | L | 5 |
| | I | 7 |
| | V | 3 |
| | M | 1 |
| | F | 5 |
| | Y | 4 |
| 804A | G | 6 |
| | P | 1 |
| | D | 156 |
| | E | 34 |
| | K | 6 |
| | S | 22 |
| | T | 10 |
| | N | 60 |
| | Q | 43 |
| | A | 23 |
| | L | 8 |
| | M | 6 |
| | F | 4 |
| | W | 1 |
| | Y | 157 |

| | | |
|---|---|---|
| 809S | S | 535 |
| | T | 1 |
| | A | 1 |
| 810Y | I | 2 |
| | V | 9 |
| | F | 107 |
| | Y | 419 |

**Table S3**: EphB1 variants selected for Michaelis-Menten analysis. Variants are named using an A-B-C-D scheme, where A refers to the design model, B refers to a run number, C refers to the sample number, and D refers to a design temperature.

| Number | ID | $k_{cat}$ ($s^{-1}$) | $K_M$ (μM) | $k_{cat}/K_M$ ($s^{-1}$ $\mu M^{-1}$) |
|---|---|---|---|---|
| 1 | mpnn-8-21462-temp0.3 | 1.36±0.01 | 53.3±14.7 | 0.025±0.007 |
| 2 | mpnn-58-3760-temp0.3 | 2.03±0.01 | 59.8±14 | 0.034±0.008 |
| 3 | f2s-71-2765-temp0.3 | 6.85±0.02 | 97.2±19 | 0.07±0.014 |
| 4 | f2s-79-7697-temp0.65 | 11.24±0.04 | 84.1±15.7 | 0.134±0.025 |
| 5 | mpnn-24-22040-temp0.3 | 1.3±0.004 | 69.7±11.8 | 0.019±0.003 |
| 6 | f2s-48-5441-temp0.3 | 3.46±0.01 | 47.5±9.6 | 0.073±0.015 |
| 7 | mpnn-45-9690-temp0.5 | 0.1±0.001 | 50.2±34.8 | 0.002±0.001 |
| 8 | mpnn-68-819-temp0.5 | 10.84±0.03 | 78.1±11.9 | 0.139±0.021 |
| 9 | mpnn-30-17043-temp0.5 | 4.17±0.01 | 38.8±6.1 | 0.107±0.017 |
| 10 | mpnn-1-22469-temp0.5 | 6.52±0.02 | 56.9±11.5 | 0.115±0.023 |
| 11 | mpnn-39-16796-temp0.5 | 13.48±0.05 | 80.6±16.4 | 0.167±0.034 |
| 12 | mpnn-39-16596-temp0.1 | 2.41±0.01 | 68.4±11.6 | 0.035±0.006 |
| 14 | mpnn-34-5159-temp0.3 | 5.44±0.02 | 40.2±9.5 | 0.135±0.032 |
| n/a | WT | 3.99±0.01 | 51.5±9.6 | 0.078±0.014 |

**Table S4:** Model predictions for unseen variants within the library. Bins are numbered according to predicted activity (1: top 8, 2: high, 3: medium, 4: low/inactive).

| ID | Predicted Yeo-Johnson Transformed Activity | Predicted Activity (RFU $min^{-1}$ $\mu M^{-1}$) | Bin | Hamming distance to closest relative in training | Number of mutations in Top10 residues | Mutations in Top 10 residues |
|---|---|---|---|---|---|---|
| mpnn-67-3329-temp0.1 | 26.3 | 831 | 4 | 7 | 8 | R662K E684L R691E D706L T791L R799G K800D S803P |
| mpnn-82-25178-temp0.5 | 15.5 | 213 | 4 | 23 | 8 | R662K E684L K689R R691E D706Y T791M R799G K800D |
| f2s-4-5740-temp0.3 | 16.2 | 239 | 4 | 10 | 8 | E684Y R691E D706K N739K T791M R799A K800T S803A |
| f2s-19-18097-temp0.3 | 24.2 | 664 | 4 | 12 | 9 | R662K E684L K689S R691S D706R N739K R799A K800T S803G |
| f2s-60-25874-temp0.3 | 30.3 | 1199 | 4 | 14 | 7 | E684L R691E D706K N739K T791M K800T S803A |

| | | | | | | |
|---|---|---|---|---|---|---|
| f2s-19-18160-temp0.65 | 31.3 | 1316 | 4 | 19 | 7 | R662K E684L K689S D706K R799A K800T S803T |
| mpnn-42-9366-temp0.3 | 58.7 | 7280 | 3 | 13 | 5 | R662K E684L R691E K800D S803P |
| f2s-36-13621-temp0.65 | 54.7 | 6006 | 3 | 15 | 5 | E684F R691E D706K K800T S803G |
| mpnn-63-17471-temp0.1 | 58 | 7058 | 3 | 9 | 7 | R662K E684L R691E D706R N739G K800D S803P |
| mpnn-59-2220-temp0.3 | 95.4 | 28540 | 2 | 11 | 7 | R662K E684L R691E D706G R799G K800D S803P |
| mpnn-12-21811-temp0.3 | 128 | 65279 | 1 | 21 | 7 | R662E E684L R691E D706R R799G K800D S803P |
| mpnn-35-2879-temp0.5 | 103 | 35630 | 1 | 22 | 7 | R662E E684V K689N R691K D706G N739G S803P |
| f2s-66-29580-temp0.65 | 67.9 | 10958 | 3 | 16 | 6 | E684Y R691N D706K N739K K800T S803G |

| | | | | | | |
|---|---|---|---|---|---|---|
| mpnn-98-15195-temp0.3 | 112 | 44743 | 1 | 8 | 7 | R662E E684L R691K D706G N739G K800N S803P |
| f2s-47-1978-temp0.65 | 85.3 | 20764 | 2 | 12 | 6 | R662A E684L R691E D706L N739E K800T |
| f2s-51-26720-temp0.3 | 93.3 | 26787 | 2 | 6 | 6 | R662K E684L R691E D706L N739K S803G |
| mpnn-39-16588-temp0.1 | 85.3 | 20764 | 2 | 4 | 7 | R662K E684L R691E D706L N739G K800D S803P |
| mpnn-40-23364-temp0.5 | 117 | 51128 | 1 | 10 | 7 | R662I E684L R691Q D706R R799G K800R S803C |
| f2s-91-12370-temp0.3 | 117 | 50991 | 1 | 7 | 7 | R662E E684L R691K D706L N739K K800T S803G |
| f2s-39-10589-temp0.3 | 129 | 47784 | 1 | 10 | 7 | R662E E684L R691E D706K N739K K800T S803K |
| f2s-54-11515-temp0.65 | 129 | 67295 | 1 | 18 | 6 | R662E E684Y R691K D706L |

| | | | | | | |
|---|---|---|---|---|---|---|
| | | | | | | K800T<br>S803K |
| f2s-77-6352-temp0.3 | 121 | 56215 | 1 | 8 | 6 | R662E<br>E684L<br>D706K<br>N739K<br>K800T<br>S803K |

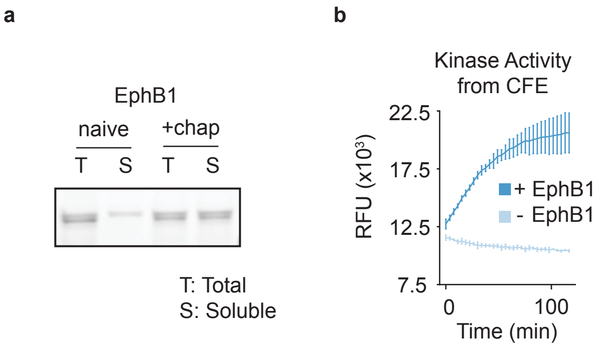


**Supplementary Figure 1**: EphB1 is solubly expressed and active in CFE reactions. (a) In-gel fluorescence images of EphB1 labelled with FluoroTect reagent during CFE show robust solubility in the presence of protein folding chaperones. (b) PhosphoSens assays are compatible with CFE reactions and can measure EphB1 activity. CFE reactions either contained linear DNA encoding EphB1 (+ EphB1) or water in place of DNA (- EphB1), and all reactions contained peptide substrate. Data shows the average of n = 3 replicates and error bars represent one standard deviation. RFU stands for relative fluorescence units.

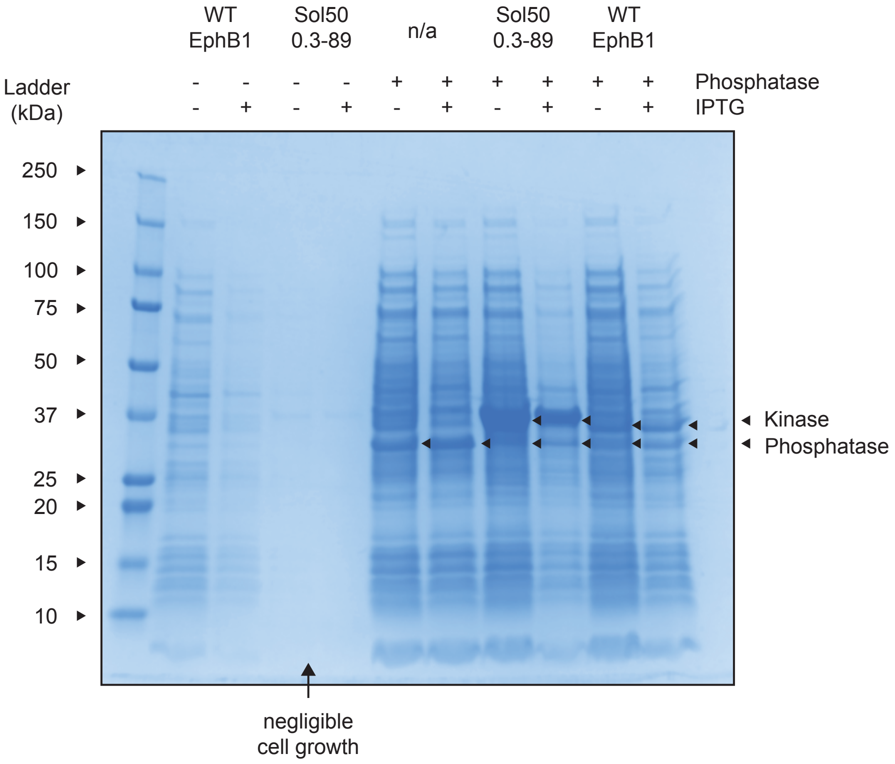


**Supplementary Figure 2:** EphB1 is toxic to *E. coli* cells in the absence of a corresponding phosphatase. Equal amounts of cell cultures overexpressing kinase or phosphatase or both were added to SDS-PAGE sample buffers, lysed by heat denaturation, and analyzed by SDS-PAGE. Sol50 0.3-89 is a redesigned EphB1 kinase which shows improved expression levels and increased toxicity (**Fig S4,5**). Arrows indicate overexpression bands of the phosphatase and the kinase.

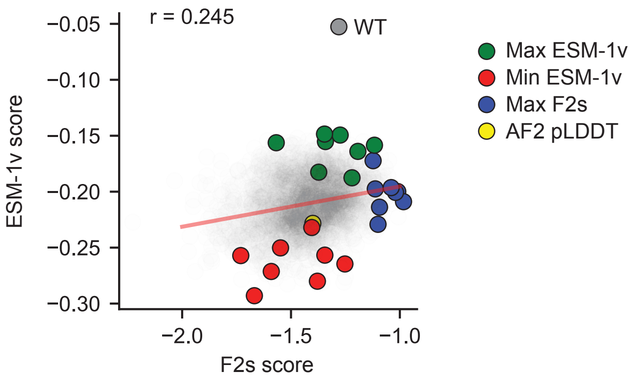


**Supplementary Figure 3**: Scatter plot of ESM-1v scores against F2s scores showing that F2s scores are only weakly correlated to ESM-1v scores. F2s scores reflect the average predicted probabilities (over all residues) that the sequence folds into a given structure, while ESM-1v scores reflect the average predicted probabilities (over all residues) from a protein language model trained on natural sequence. The highlighted points represent sequences chosen for experimental testing, as well as the wild-type sequence. The red line depicts the linear fit between F2s scores and ESM-1v sequences (Pearson r = 0.245).

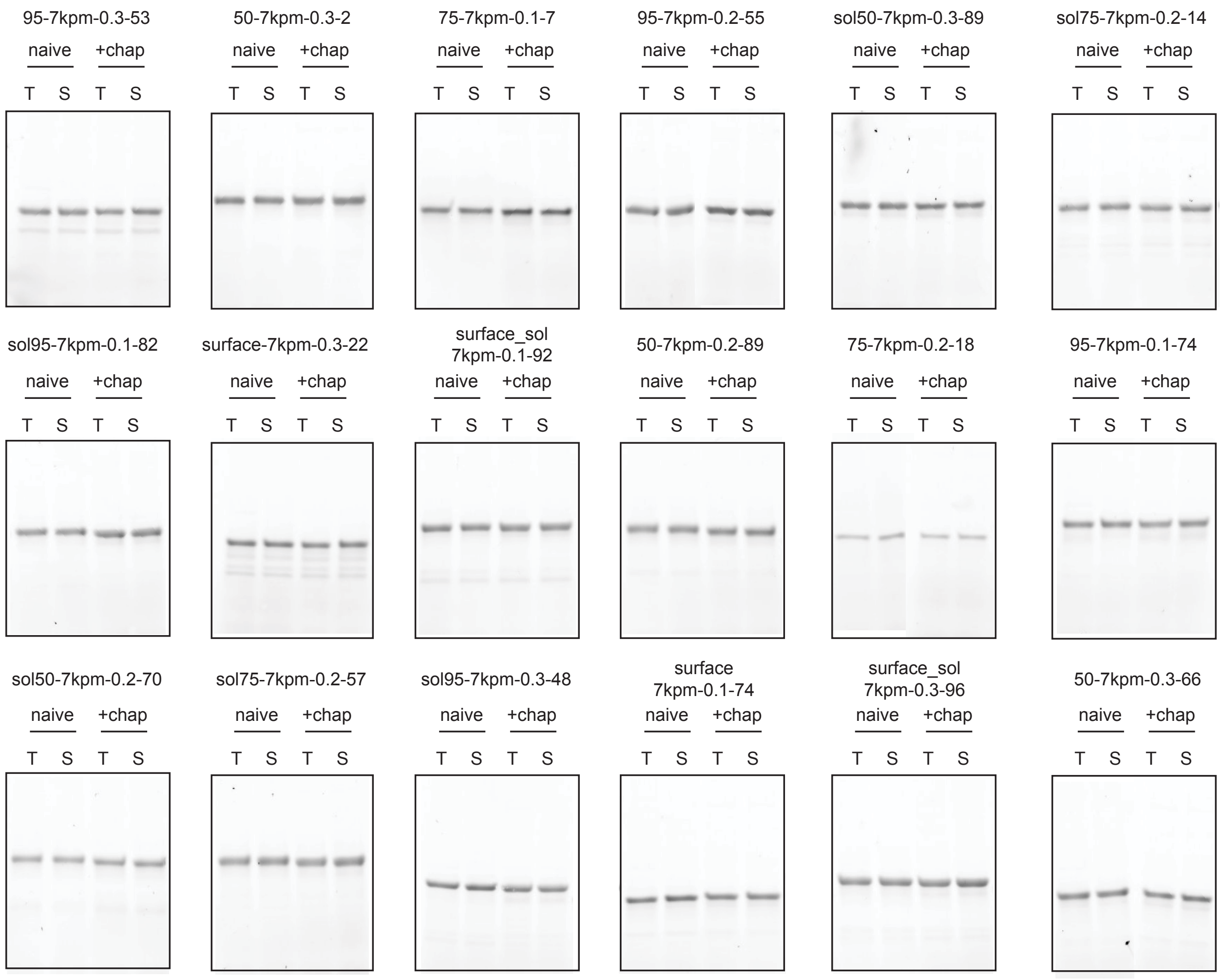


Additional variants on next page...

**Supplementary Figure 4** (continued on next page)

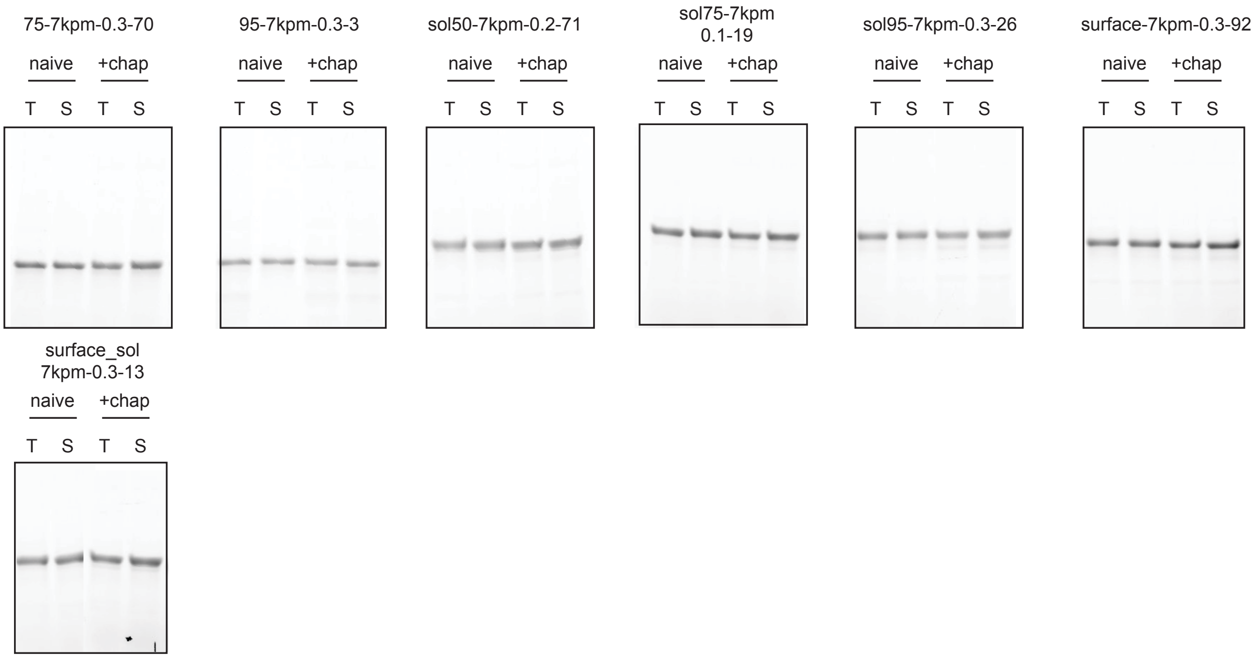


**Supplementary Figure 4:** FluoroTect gels showing that redesigned EphB1 variants express well even in the absence of protein folding chaperones. Band positions vary between redesigned sequences, which is likely due to increased stability after pMPNN redesign or changes in charge (redesigned sequences are often highly charged). All gels were run simultaneously. Images for 95-7kpm-0.2-55, 75-7kpm-0.2-18, 50-7kpm-0.3-66, surface_sol-7kpm-0.3-13 are consolidated from two separate gels. Raw gels are available as supplementary files.

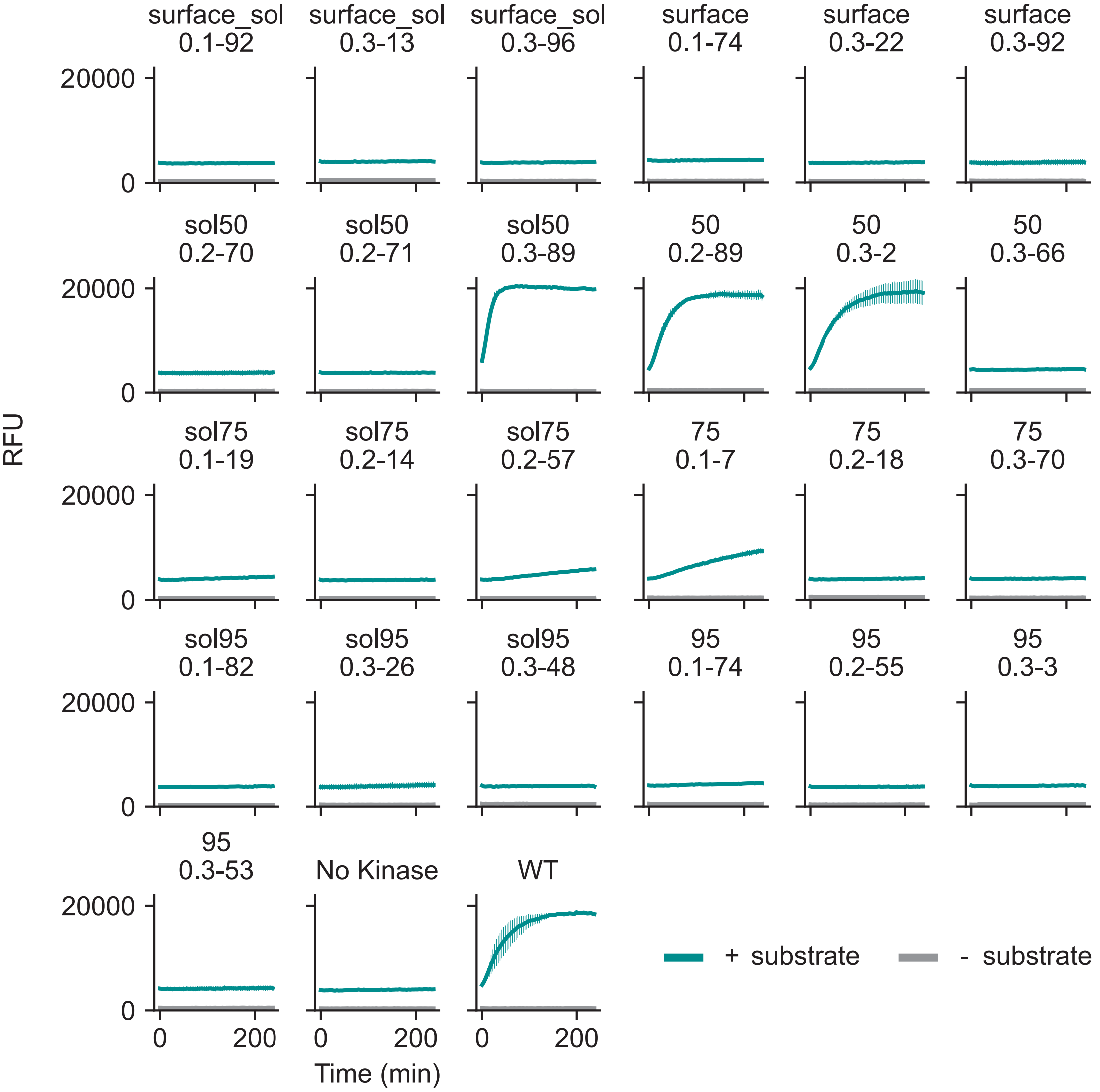


**Supplementary Figure 5**: Five redesigned EphB2 variants from the pilot library are active in PhosphoSens assays. Plots show fluorescence (measured in relative fluorescence units (RFU)) over time in the presence (in green) or absence (in grey) of a substrate with a given kinase variant. The best variant, Sol50-0.3-89, shows improved activity over WT likely due to higher expression levels (**Fig S2**). Solid line represents the mean over n = 3 replicates with error bars representing one standard deviation.

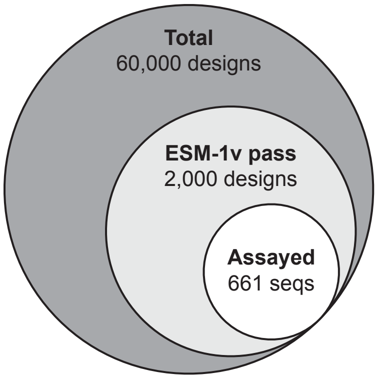


**Supplementary Figure 6**: Schematic for the sequence design and filtering pipeline in the second generation EphB1 library.

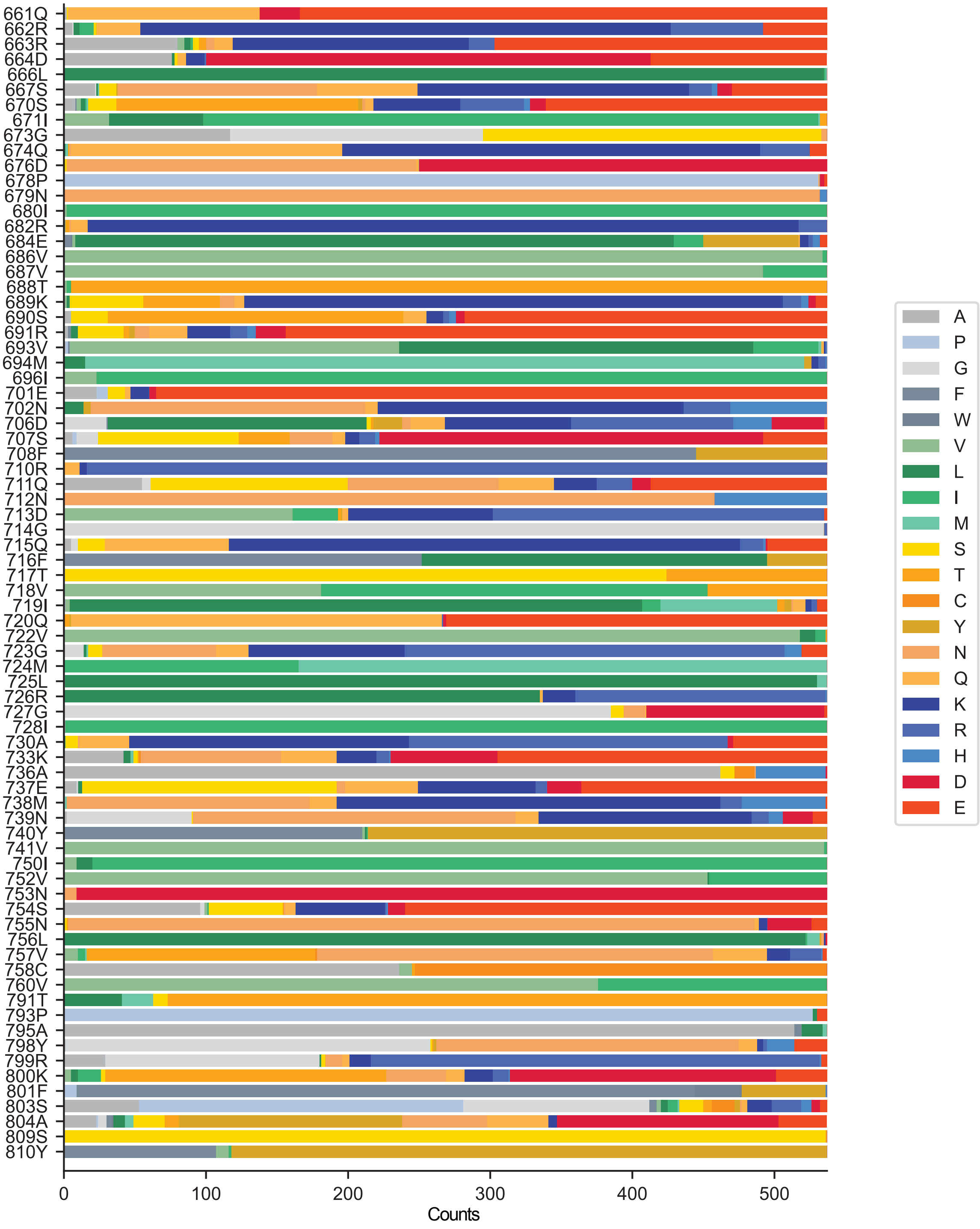


**Supplementary Figure 7**: Bar plot showing counts of all mutations at all redesigned positions in the redesigned EphB1 variants tested in our workflow.

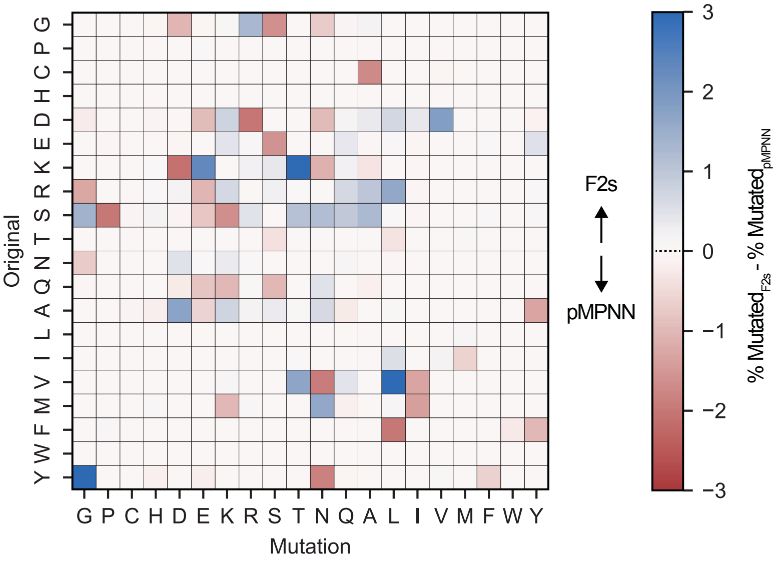


**Supplementary Figure 8**: The difference matrix between substitution matrices for F2s and pMPNN sequences highlights mutational preferences for each model.

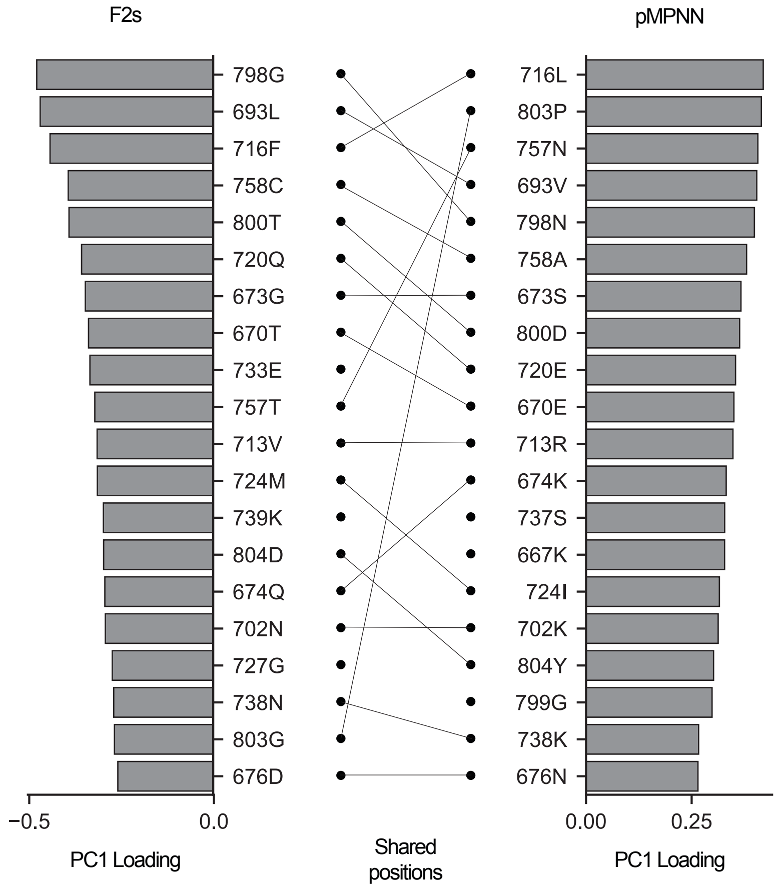


**Supplementary Figure 9**: The top 20 loadings onto PC1 in a principal component analysis reveal mutational signatures between sequences designed by F2s and pMPNN. Lines connecting two positions indicate that the models redesign the same position with different amino acids.

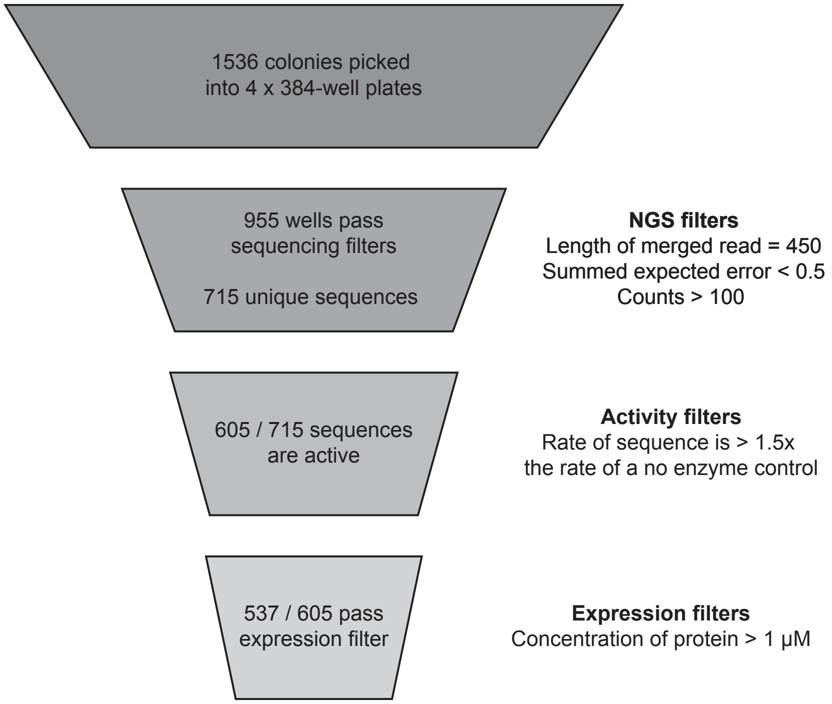


**Supplementary Figure 10**: Filtering pipeline for high quality sequence-function measurements.

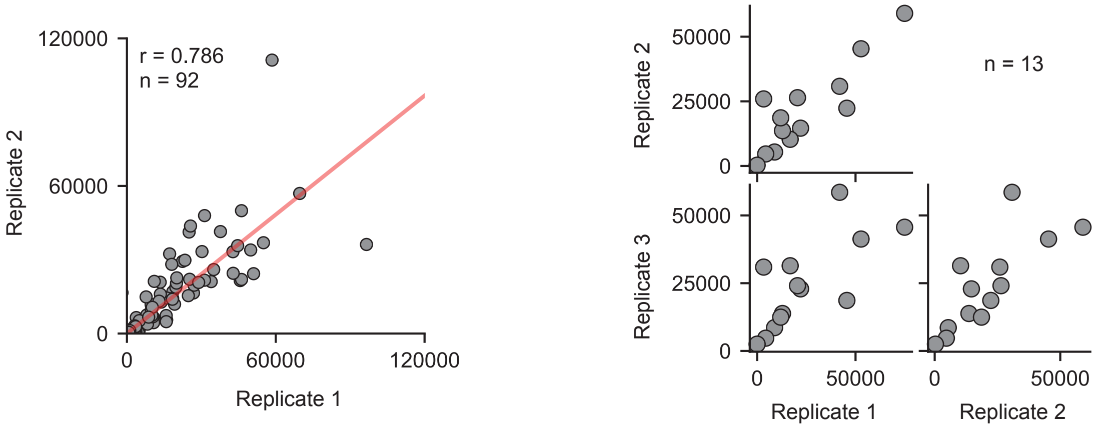


**Supplementary Figure 11**: Internal replicates of specific activity measurements within the dataset are highly correlated. Data show replicates between separate measurements of Specific Activity (RFU $min^{-1}$ $\mu M^{-1}$) on the same sequence. Duplicates show consistency between measurements (Pearson r = 0.786).

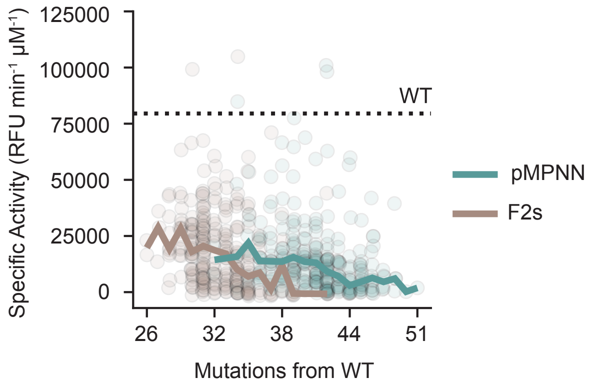


**Supplementary Figure 12:** Specific activity decreases with increasing number of mutations from the WT. Solid line indicates the median of specific activities at each number of mutations. Each point represents a single sequence and its associated hamming distance from WT and Specific Activity measurement. Color represents whether the sequence was designed by pMPNN (teal) or F2s (brown).

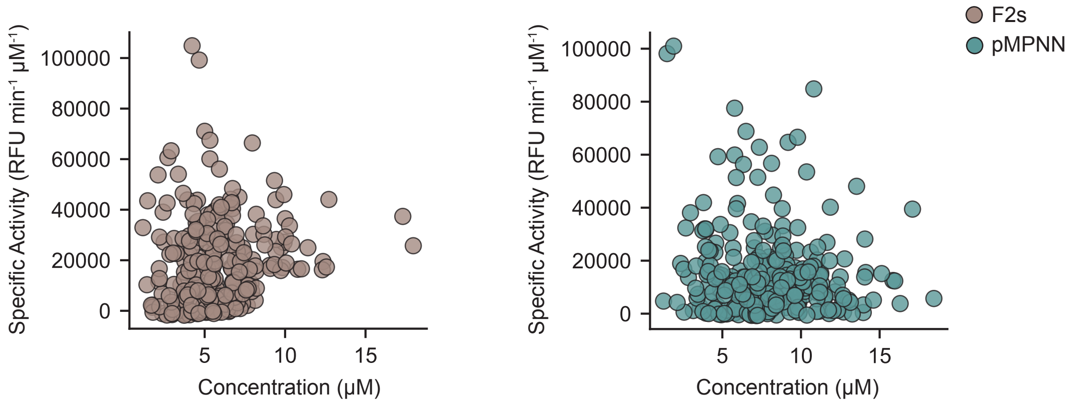


**Supplementary Figure 13:** Expression and activity tradeoffs in F2s and pMPNN sequences. Each datapoint represents a specific sequence and its associated Specific Activity and expression levels.

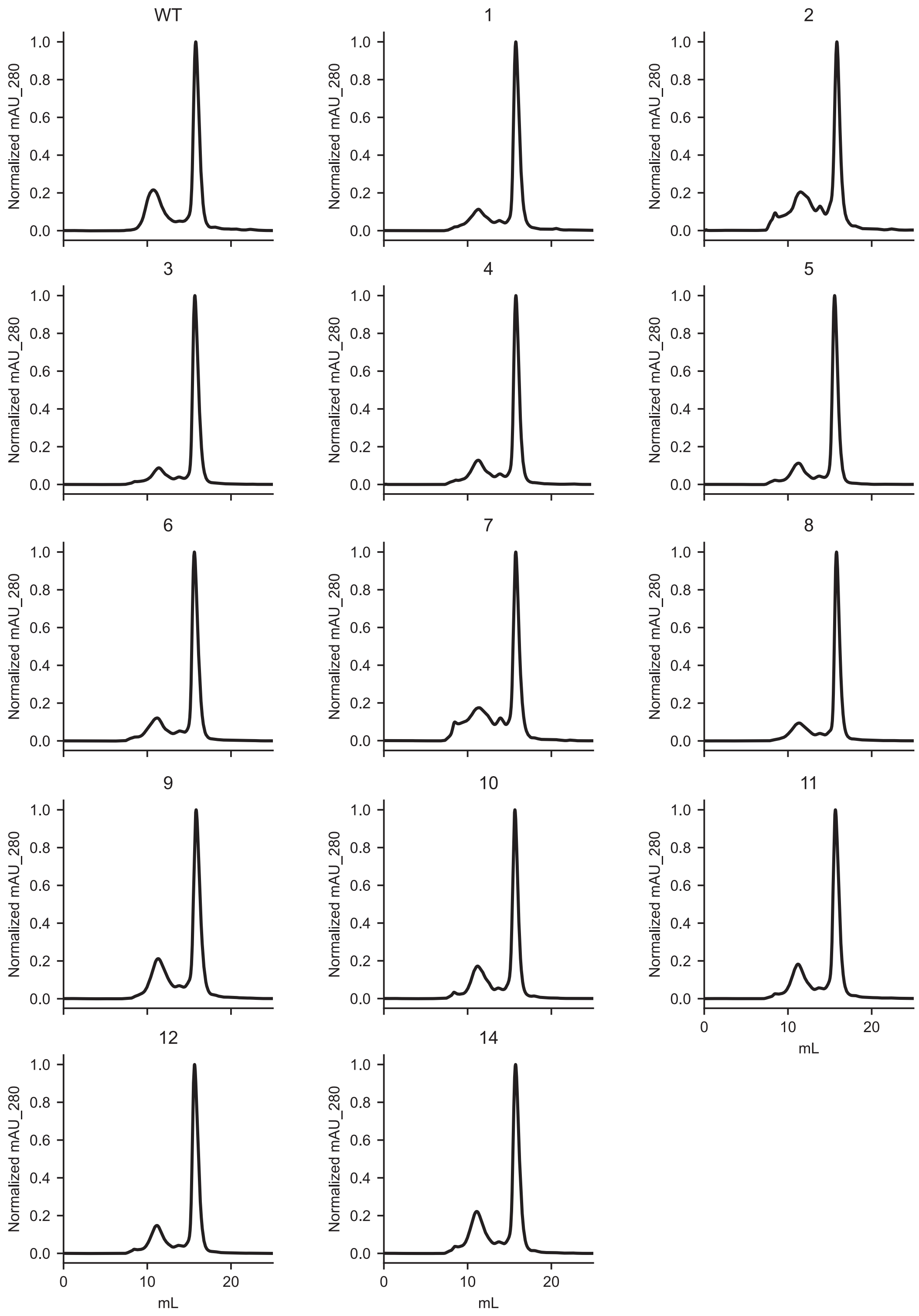


**Supplementary Figure 14**: Redesigned EphB1 run essentially identical to the WT in size-exclusion chromatography. Only the monomeric peak appearing at column volumes of 14-17 mL was collected for Michaelis-Menten analysis. Numbers refer to variants as labelled in **Table S2**.

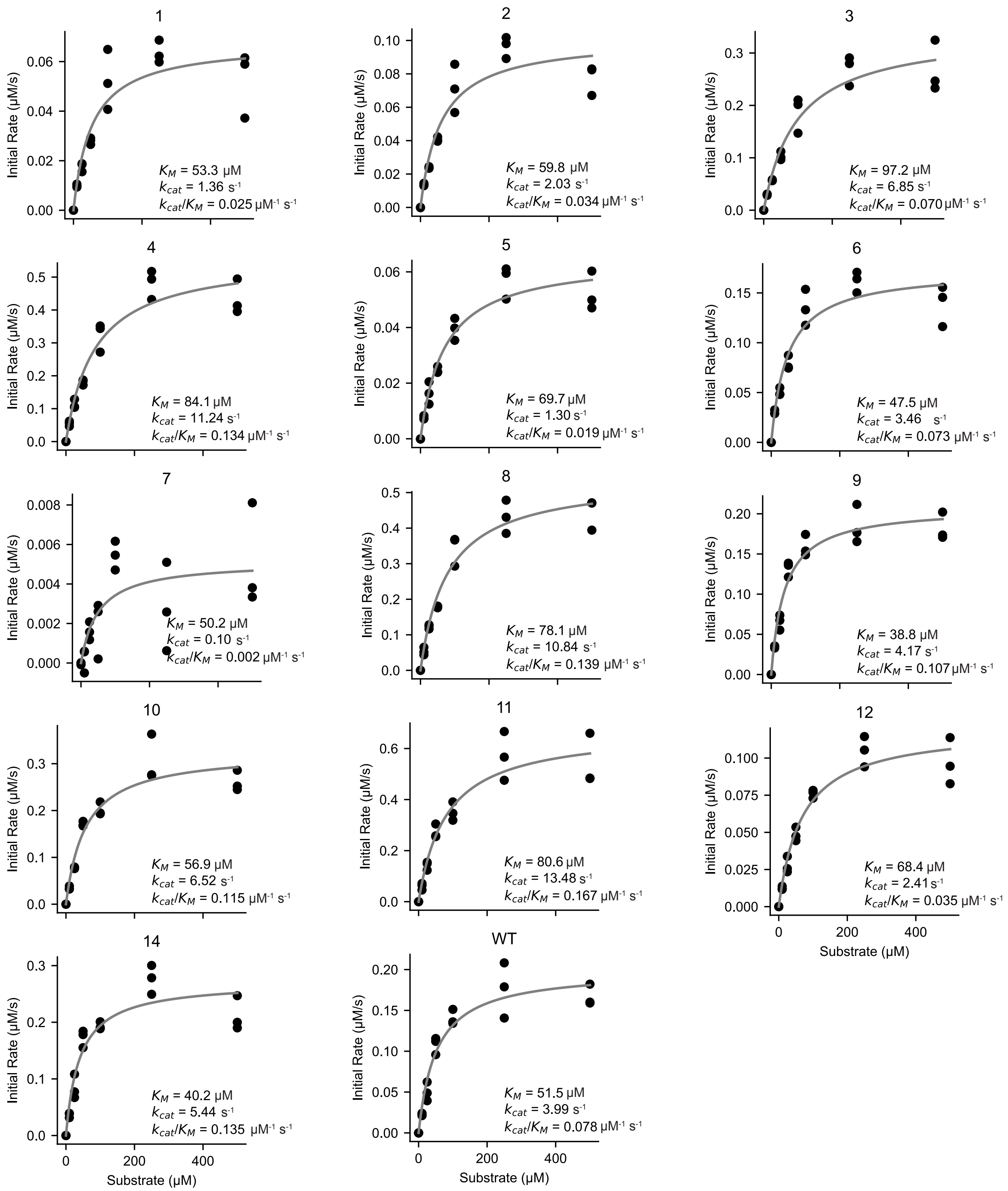


**Supplementary Figure 15**: Michaelis-Menten analysis of 13 redesigned EphB1 kinase variants. Numbers refer to variants as labelled in **Table S2.** Gray lines represent best fit line to Michaelis-Menten equation:

$$Initial\ Rate = \frac{V_{max}[S]}{K_M+[S]} \text{ where } V_{max} = [E]_{total}k_{cat}$$

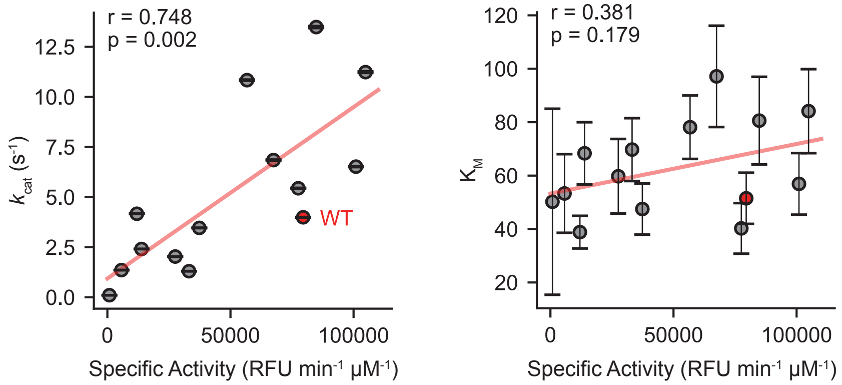


**Supplementary Figure 16:** $k_{cat}$ measurements for redesigned EphB1 variants correlate well with Specific Activity measurements. $K_M$ measurements do not correlate strongly with activity. Points represent the average of n = 3 replicates and error bars represent one standard deviation of the fits. Red line denotes the line of best fit ($k_{cat}$: Pearson r = 0.748, p = 0.002 (Wald test), $K_M$: Pearson r = 0.381, p = 0.719 (Wald test)).

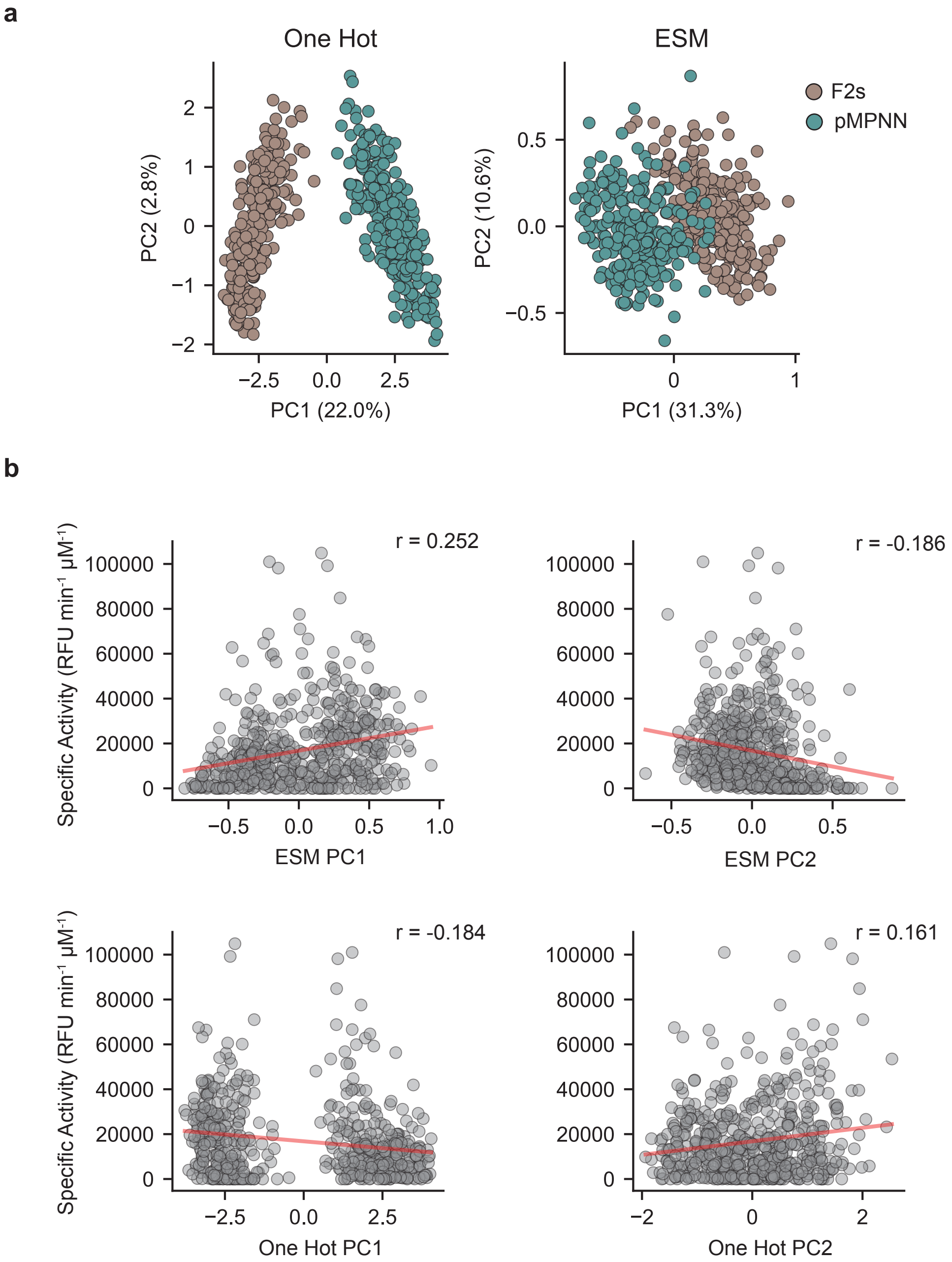


**Supplementary Figure 17**: Principal component analysis does not cluster sequences by activity level. (**a**) PCA of one hot encoded sequences or of ESM-1v embeddings of sequences separates by design algorithm along PC1. (**b**). Principal components do not correlate well with activity. Red lines represent lines of best fit with Pearson r values (ESM PC1: r = 0.252, ESM PC2: r = -0.186, One Hot PC1: r = -0.184, One Hot PC2 r = 0.161).

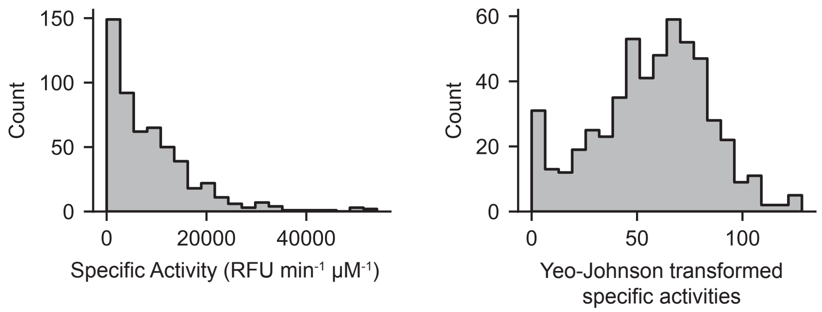


**Supplementary Figure 18**: Specific activities were transformed towards normality using the Yeo-Johnson transformation. Distributions of specific activities and transformed values are shown as histograms.

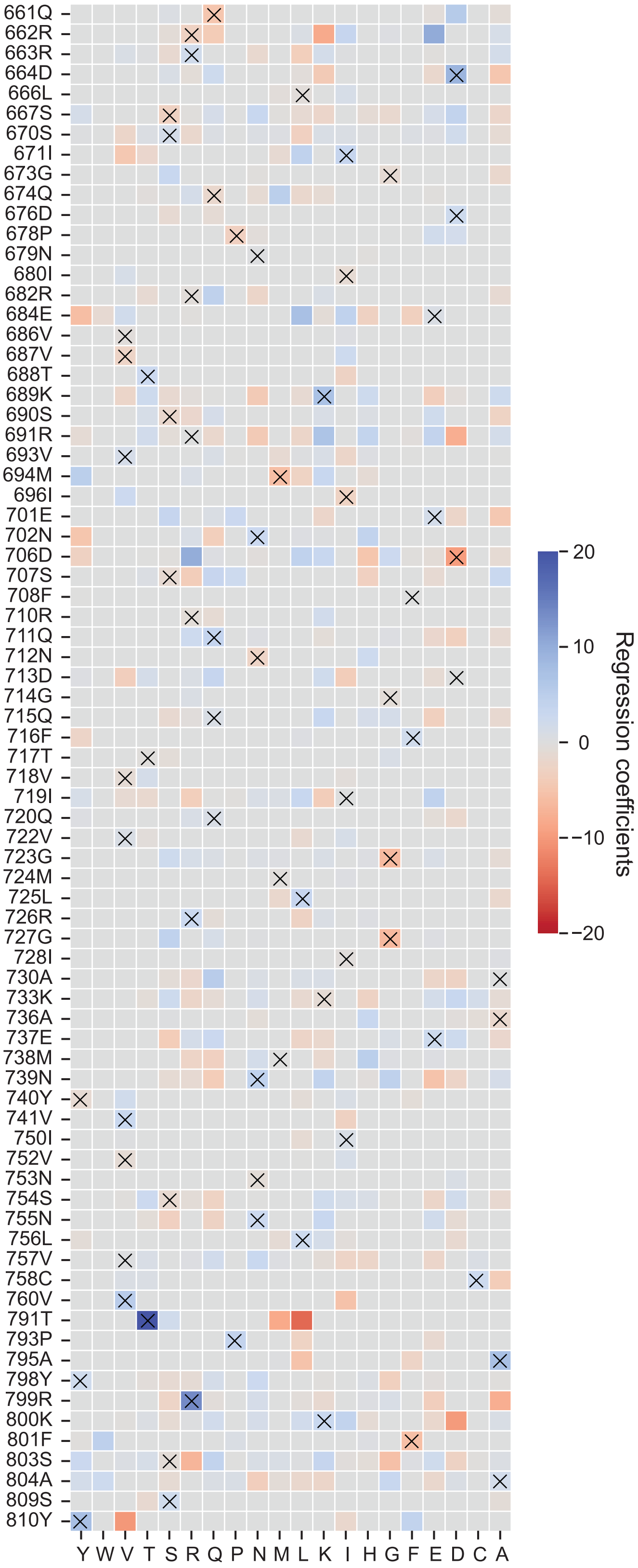


**Supplementary Figure 19**: Regression coefficients for the 20 amino acids (one-letter code) of ridge regression model. X in the heatmap denotes the WT residue at that position.

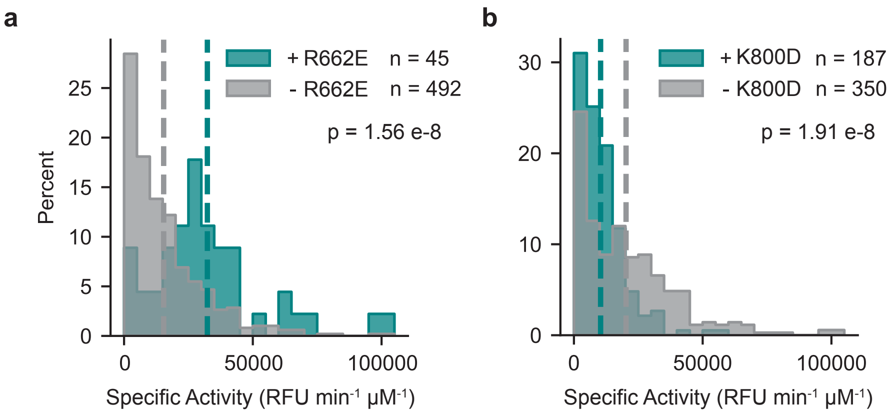


**Supplementary Figure 20**: Histograms of sequence Specific Activities in the presence and absence of (**a**) R662E or (**b**) K800D. p values (1.56e-8 and 1.91e-8 for **a** and **b**, respectively) are calculated by Mann-Whitney U test

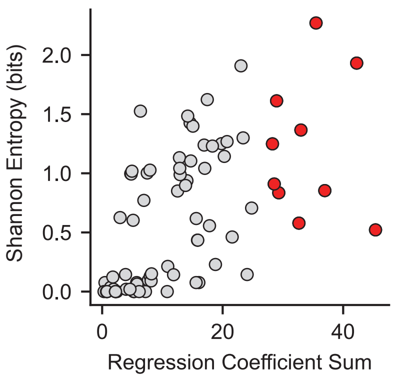


**Supplementary Figure 21**: Positions that contribute most to model performance (red) have widely different Shannon entropies. Mutations at every designed position were grouped into physiochemical categories (Hydrophobic: A,L,I,V,M; Aromatic: F,W,Y; Polar: S,T,N,Q; Charged: D,E,K,R; Glycine: G, Proline: P, Cysteine: C, Histidine: H) and Shannon entropy was calculated for each position based on these categories.

**a**

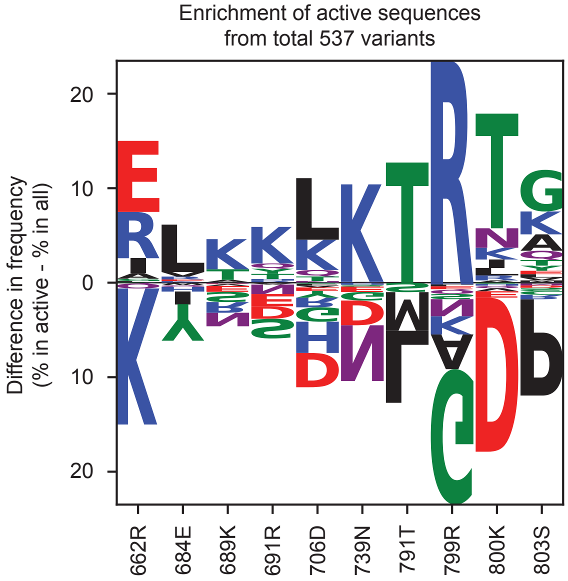


**b**

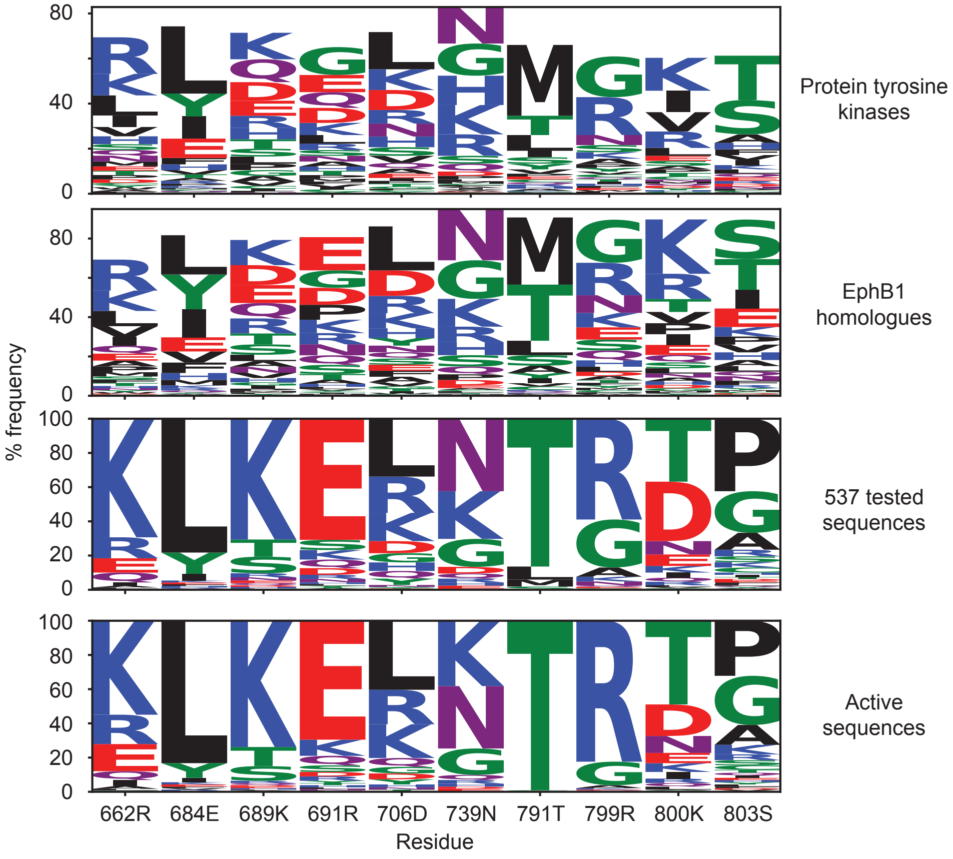


**Supplementary Figure 22**: Weblogos of mutation frequences in the top 10 positions identified by a ridge regression model in kinases. (**a**) Enrichment of mutations in redesigned EphB1 kinases at the top 10 positions in 537 experimentally tested sequences and in active sequences (> 15,000 RFU $min^{-1}$ $\mu M^{-1}$). (**b**) Frequency of mutations in the top 10 positions in representative protein tyrosine kinases, in Eph tyrosine kinase homologues, in 537 experimentally tested sequences, and in active variants.

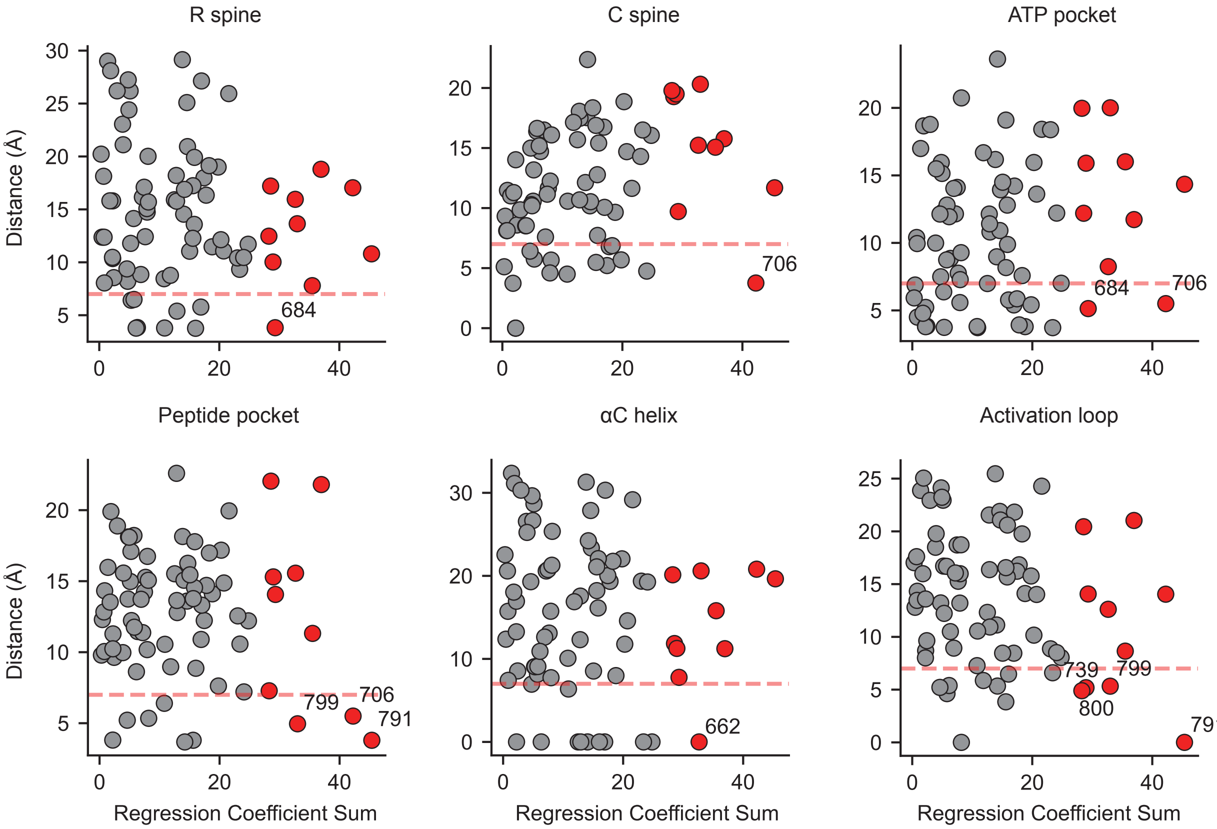


**Supplementary Figure 23**: Scatterplots showing the relationship between residue importance (as measured by sum of ridge regression coefficients) to distance to functional regions of protein kinases. Distance is defined as the shortest distance between Cα atoms at a redesigned position and a residue position in a functional region (denoted in plot title). Seven of the top 10 residues are within a Cα distance < 7 Å to a residue within a functional region. Points in grey indicate residue positions that were redesigned. The top 10 most important residues are labelled in red. Dotted red line indicates Cα distance = 7 Å.

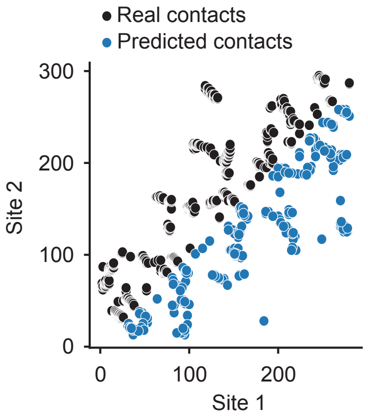


**Supplementary Figure 24**: Potts model, in blue points, captures real contacts in the EphB1 structure, in black points. Contacts are defined as residues whose $C_\alpha$ are within 8Å.
