## Supplementary figures and images for "A combinatorial mutational map of active non-native protein kinases by deep learning guided sequence design"

### Raw uncropped gels

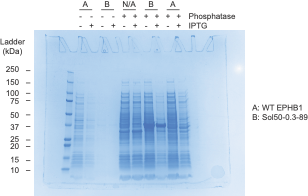

**Raw gel for Supplementary Figure 2**

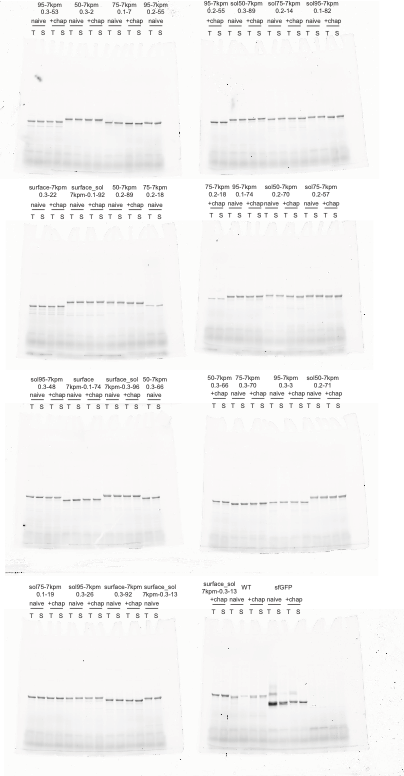

**Raw gel images for Supplementary Figure 4**
